# Mapping the adaptor protein complex interaction network in *Arabidopsis* identifies P34 as a common stability regulator

**DOI:** 10.1101/2022.08.31.505729

**Authors:** Peng Wang, Wei Siao, Xiuyang Zhao, Deepanksha Arora, Ren Wang, Dominique Eeckhout, Jelle Van Leene, Rahul Kumar, Anaxi Houbaert, Nancy De Winne, Evelien Mylle, Michael Vandorpe, Ruud A. Korver, Christa Testerink, Kris Gevaert, Steffen Vanneste, Geert De Jaeger, Daniël Van Damme, Eugenia Russinova

## Abstract

Adaptor protein (AP) complexes are evolutionarily conserved vesicle transport regulators that recruit coat proteins, membrane cargos and coated vesicle accessory proteins. Since in plants endocytic and post-Golgi trafficking intersect at the *trans*-Golgi network, unique mechanisms for sorting cargos of overlapping vesicular routes are anticipated. The plant AP complexes are part of the sorting machinery, but despite some functional information, their cargoes, accessory proteins, and regulation remain largely unknown. Here, by means of various proteomics approaches, we generated the overall interactome of the five AP and the TPLATE complexes in *Arabidopsis thaliana*. The interactome converged on a number of hub proteins, including the thus far unknown adaptin binding-like protein, designated P34. P34 interacted with the clathrin-associated AP complexes, controlled their stability and, subsequently, influenced clathrin-mediated endocytosis and various post-Golgi trafficking routes. Altogether, the AP interactome network offers substantial resources for further discoveries of unknown endomembrane trafficking regulators in plant cells.

## Introduction

The transport of integral membrane proteins from one compartment to another subcellular destination within the endomembrane system is essential for all living eukaryotic cells. This transport is mostly mediated by coated vesicles that are formed by the recruitment of coat proteins onto a particular membrane compartment where they bind adaptors, and other accessory molecules to promote membrane curvature and collect the vesicle cargo^1^. Critical for this process is the fine-tuned control of intracellular cargo sorting, which relies on signals within the cargo proteins and on the specific adaptor proteins (AP) complexes-mediated cargo recognition. Five AP complexes, designated AP-1 to AP-5, each involved in a distinct vesicle trafficking pathway have been identified in higher eukaryotes^2^. The five AP complexes are heterotetrameric proteins that contain two large subunits (β1 to β5 and γ, α, δ, ε, ζ), a medium subunit (µ1 to µ5) and a small subunit (σ1 to σ5)^3^. Whereas in mammalian cells, AP-1, AP-2 and AP-3 operate in concert with the scoffing molecule clathrin to generate clathrin-coated vesicles (CCVs), AP-4 and AP-5 function independently of clathrin^3^.

In plant cells, the major membrane trafficking, including secretory, endocytic and vacuolar pathways intersect at the *trans*-Golgi network (TGN)/early endosome (EE)^4^. Over the last decade, some functions of AP-1 to AP-4 have been characterized in the model plant *Arabidopsis thaliana*^5^. Although, sequence-based homology searches have predicted a partial AP-5 in plant genomes^2^, the AP-5 function and composition remain undetermined^6^. Plant AP-1 associates with TGN/EE and transports cargos to the plasma membrane (PM), cell plates or vacuoles, and loss of the AP-1 function leads to impeded growth, cytokinesis defects, and male and female sterility^7–10^. Plant AP-2 facilitates clathrin-mediated endocytosis (CME) of various PM proteins, such as the PIN-FORMED^11^ proteins, BR INSENSITIVE1 (BRI1)^12, 13^ receptor and the boron transporter REQUIRES HIGH BORON1 (BOR1)^14^, and is important for reproductive organ development^15, 16^. AP-3 enables the vacuolar targeting of the SUCROSE TRANSPORTER4 (SUC4) and the R-SNARE subfamily members VAMP713, and loss of AP-3 function reduces germination and affects gravitropic responses^17–20^. AP-4 mediates vacuolar protein sorting and the recruitment of the ENTH domain protein, MODIFIED TRANSPORT TO THE VACUOLE1 (MTV1) at the TGN/EE, and AP-4 deficiency causes growth defects and abolishes hypersensitive cell death upon infection with avirulent bacteria^21–24^. Alongside AP-2, an ancient heterooctameric adaptor complex, designated the TPLATE complex (TPC), also is required for CME at PM in plant cells^25, 26^. Knockout of single subunits of TPC caused male sterility, whereas silencing triggered seedling lethality^25, 27^ and abolished the endocytic flux, as also observed upon conditional inactivation of the complex^28^.

In addition to the coat proteins and the tetrameric AP complexes, CME in mammalian and yeast cells requires the activity of more than 60 endocytic accessory proteins (EAPs) that function as scaffolds, cargo recruiters, membrane curvature generators and sensors, as well as CME regulators^29, 30^. Such EAPs have been found either by comparative proteomics of CCVs^31, 32^ or by affinity purification coupled to mass spectrometry (AP-MS) as binding partners of the C-terminal appendages or EAR domains of the large subunits of AP complexes^33, 34^. Although, some EAPs have been identified based on sequence homology^4, 6^ and recently by proteomic analysis of Arabidopsis CCVs^35^, their involvement in vesicle formation in plant cells remains largely uncharacterized.

Here, using AP-MS and proximity labelling (PL) coupled to MS (PL-MS), we generated a comprehensive interactome of the six adaptor protein complexes in Arabidopsis, thereby providing informative resources and links between vesicle transports and their regulators. We identified the AP-5 complex and its associated core subunits, spatacsin (SPG11) and spastizin (SPG15 also known as ZFYVE26), in Arabidopsis cells. The interactomes of the clathrin-associated AP complexes in Arabidopsis, AP-1, AP-2 and AP-4^35^, converged on several hub proteins, of which the adaptin binding-like protein, designated P34, was functionally characterized. The *P34* gene is essential for Arabidopsis because a loss-of-function mutation led to embryo lethality. We showed that P34 controls the stability of the AP-1, AP-2 and AP-4, but not of AP-3 and as such, influences CME and various post-Golgi trafficking routes.

## Results

### Comprehensive mapping of the AP interactomes and accessory proteins reveals two tightly interconnected networks

To gain further insight into the function of the five AP complexes and TPC in plants, we performed a targeted interaction screen in Arabidopsis cell suspension cultures (PSB-D). Hereto, we applied a series of tandem affinity purification (TAP), single-step AP-MS and PL-MS experiments (see Methods for more details). Previously, those interactome analyses were successfully used for the identification of the core AP-2 and TPC in Arabidopsis^12, 25, 36^. Upon integrating all interactome analyses including further network extension through a selection of reciprocal (T)AP-MS experiments (see below), we generated a comprehensive network of 926 interactions among 536 proteins (Fig. 1 and Supplementary Data 1).

**Fig. 1.**
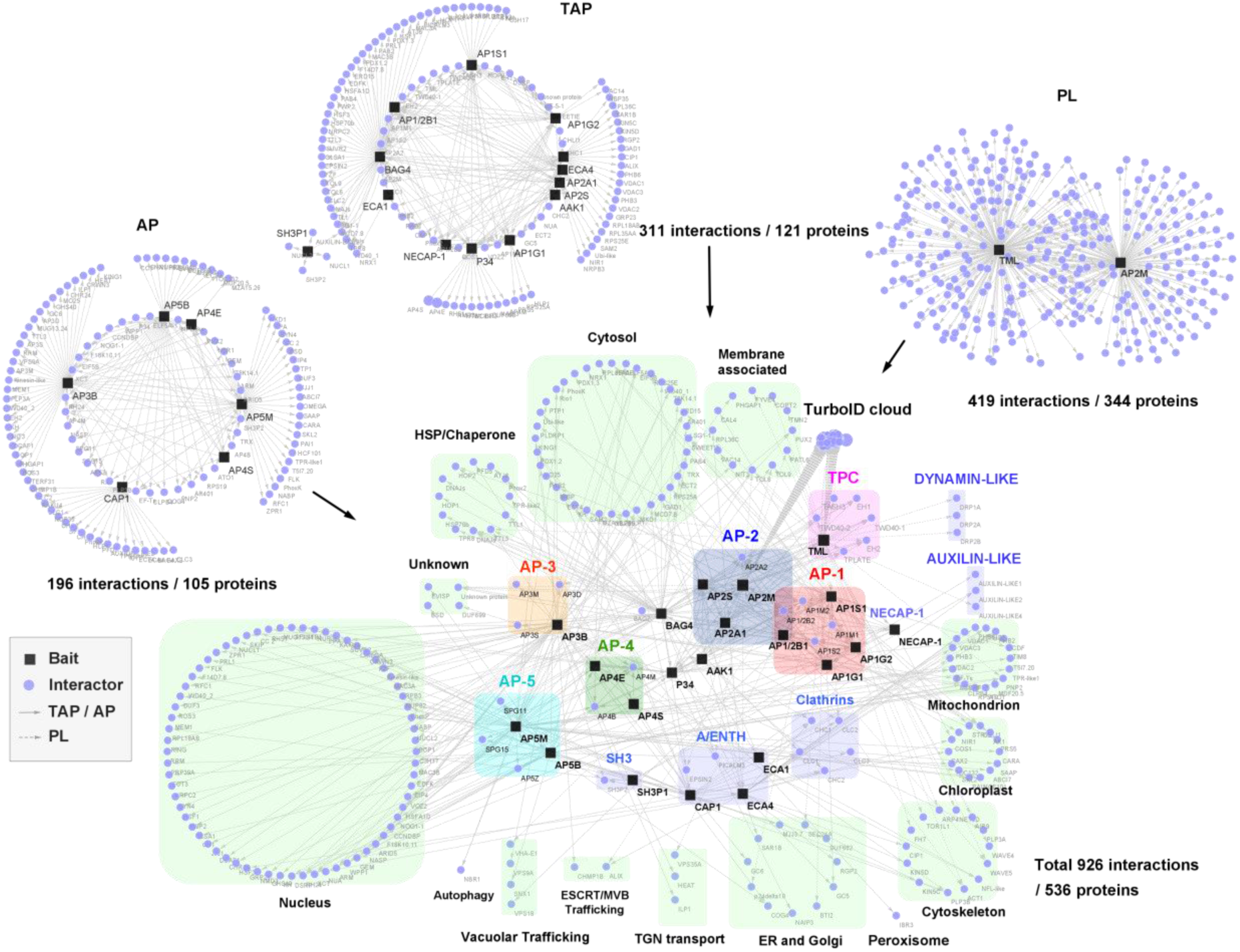
Integrative interaction network of six multimeric adaptor complexes and their accessory proteins. Thirteen proteins (AP1G1, AP1G2, AP1S1, AP2A1, AP2B1, AP2S, BAG4, P34, AAK1, NECAP-1, AtECA1, AtECA4 and SH3P1) were used as baits for TAP-MS, six proteins (AP3B, AP4E, AP4S, AP5B, AP5M and CAP1) for AP-MS, and two proteins (AP2M and TML) for PL-MS. In summary, 121 proteins with 311 interactions, 105 proteins with 196 interactions and 344 proteins with 419 interactions were captured by TAP-MS, AP-MS and PL-MS, respectively. Rounded rectangles in different colours represent the components of different complexes, including AP-1 (red), AP-2 (blue), AP-3 (orange), AP-4 (green), AP-5 (cyan), TPC (magenta) and other accessory proteins (light purple). The PL cloud indicates a group of proteins identified by PL-MS without direct link to vesicle trafficking. The proteins copurified with the baits are annotated by their known or predicted subcellular localizations as indicated (light green rectangles).

First, we analysed AP-1 and AP-2 using AP1G1, AP1G2, AP1S1, AP1/2B1, AP2A1 and AP2S as baits for TAP-MS, retrieving 110 interactions among 52 proteins (Fig. 1 and Supplementary Data 1). As a complementary approach, we mapped proximal interactors of the AP-2 and the TPC with PL using AP2M and the TPC subunit TML as bait proteins fused to the biotin ligase TurboID tag, revealing 419 interactions among 344 proteins (Fig. 1 and Supplemental Data 1). Both the TAP and PL approaches were successful, because all the core subunits of AP-1, AP-2 and TPC (except for the smaller LOLITA protein) were identified (Extended Data Fig. 1). Consistent with the fact that AP-2 functions in endocytosis and shares the β subunit with AP-1, AP-2 was found to associate both with the AP-1 and TPC (Fig. 2), because AP1M2, AP1S1 and all TPC subunits, except for the LOLITA, were found with AP-2 subunits as baits (Fig. 2, right inset). When the AP-1 subunits were used as baits, all AP-2 subunits, except AP2A1, were identified in these experiments, demonstrating the interconnection between AP-1 and AP-2. Besides the core AP-1, AP-2 and TPC subunits, several known accessory proteins linked to endocytosis and vesicle trafficking were detected through PL, such as such as the clathrin light chain subunit 2 (CLC2), four dynamin-related proteins (DRP1A, DRP1E, DRP2A, and DRP2B), the clathrin uncouting factors AUXILIN-LIKE1 and AUXILIN-LIKE4^37^, the R-SNARE protein VESICLE-ASSOCIATED MEMBRANE PROTEIN 721 (VAMP721)^38^, and six ENTH/ANTH/VHS accessory proteins (ECA4, PICALM3, EPSIN2, EPSIN3, TOL6, TOL9)^39^. Their specific PL identification concurs with the fact that PL enhances the detection of membrane-related protein interactions, when compared to protein complex analysis by (T)AP-MS^36^. Nonetheless, the TAP-MS analysis on the core AP-1, AP-2 and TPC subunits retrieved also additional proteins that can be linked to endocytosis, as witnessed by the co-purification of an Arabidopsis homolog (AT3G58600) of the human ADAPTIN EAR-BINDING COAT-ASSOCIATED PROTEIN-1 (NECAP-1)^40^ with three AP-1 subunits in TAP-MS. The interaction between NECAP-1 and AP-1 was confirmed in reciprocal TAP-MS experiments with NECAP-1 as a bait, in which all AP-1 subunits but none of the AP-2 subunits were detected (Fig. 1 and Supplementary Data 1). These TAP and PL analyses clearly demonstrate the robustness of the interactome data, not only providing valuable insight into the functioning of the core AP-1, AP-2, and TPC, but potentially also revealing unknown accessory proteins. Notably, three proteins with previously undetermined functions in plant endomembrane trafficking, a GTP-binding protein (AT5G65960), a protein kinase superfamily protein (AT2G32850) and the BCL-2-ASSOCIATED ATHANOGENE4 (BAG4) protein^41^ (AT3G51780), were identified with multiple AP-2 subunits and TML as bait proteins, highlighting them as novel candidate accessory proteins. Sequence homology analysis revealed that the GTP-binding protein is a putative plant homolog of the human α- and γ-adaptin binding protein (AAGAB) P34^42, 43^, with 23% identity and 38% similarity (Extended Data Fig. 1), whereas the uncharacterized protein kinase is a plant homolog of the human AP2-ASSOCIATED PROTEIN KINASE1 (AAK1) ^44, 45^, with 43% identity and 62% similarity (Extended Data Fig. 2). The BAG family protein BAG4 is a member of an evolutionarily conserved group of co-chaperones found in yeasts, animals and plants, linked mainly to apoptosis^41^. To further consolidate the predicted role of P34, BAG4 and AAK1 in plant vesicle trafficking based on sequence similarity to their mammalian orthologues^41, 42, 45^, reciprocal TAP-MS analyses using these three proteins as baits identified 87 interactions among 73 proteins (Fig. 1 and Supplemental Data 1). The reciprocal TAP-MS analyses indicated that also in plants, P34, AAK1 and BAG4 are linked with vesicle trafficking, because numerous AP-1, AP-2, AP-4, and TPC subunits, clathrin, and ENTH/ANTH/VHS proteins were retrieved. The TAP-MS experiments with AAK1 confirmed its interaction with AP-2 and clathrin heavy chains (CHCs). Intriguingly, highly stable and stoichiometric interactions were found between P34 and multiple AP-2 and AP-1 subunits, while also interacting with the AP-4 subunit, AP4M although this interaction was less prevalent. Also with BAG4, multiple members of both the AP-2 and AP-1 were co-purified, in addition to five subunits of the TPC, two CLCs and multiple ENTH/ANTH/VHS proteins. To further extend our endocytosis-related network, the three AP180-amino-terminal-homology (ANTH)-domain containing proteins ECA1, ECA4 and CAP1, which were isolated with BAG4 as a bait and are involved in membrane budding processes^39^, were selected for reciprocal TAP-MS (ECA1 and ECA4) or AP-MS (CAP1). These analyses support that ECA4 and CAP1 function is associated with CCV, as CLCs and CHCs were isolated with these baits. Moreover, ECA4 and CAP1 seem to function in one protein complex, as both proteins were abundantly co-purified which each other. Similarly, also P34, AAK1 and BAG4 were isolated with each other. In summary, this first set of interactome analyses implies that AP-1, AP-2 and TPC are intrinsically connected to each other, and that P34, BAG4 and AAK1 function as accessory proteins that link these complexes.

**Fig. 2.**
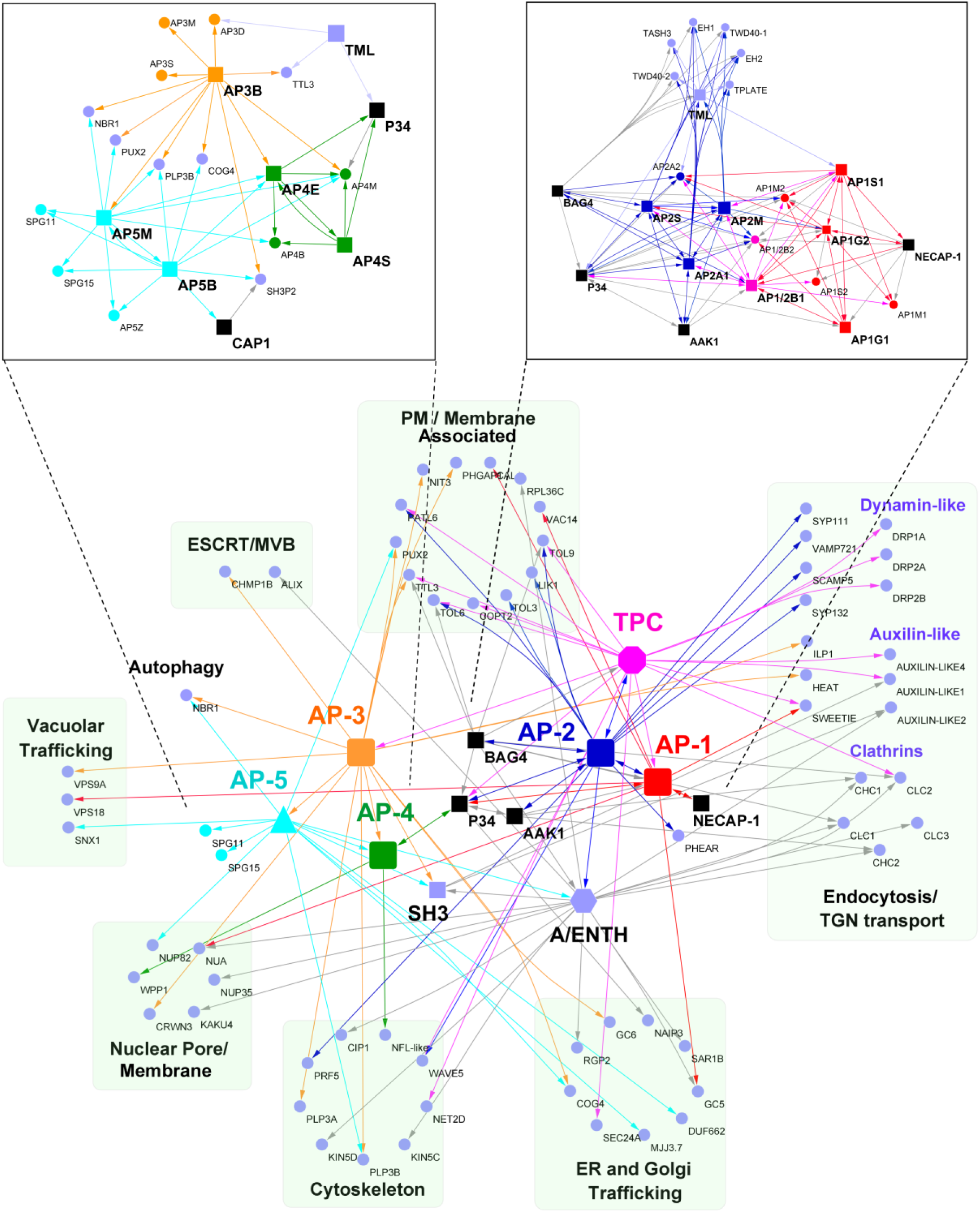
Selection from the AP interactome of putative regulators and their crosstalk in vesicle trafficking. Interactors related to membrane and protein transport were selected. The subunits of each AP complex are combined and shown as individual units (nodes) in colours (AP-1, red; AP-2, blue; AP-3, orange; AP-4, green; AP-5, cyan; TPC, magenta). The purple hexagon and square nodes represent the protein members of A/ENTH and SH3, respectively. P34, BAG4 and AAK1 proteins are shown in black squares. The purple round nodes are the interactors that are functionally linked with protein transport. Source and target nodes of the arrows indicate the baits and the targets, respectively. *Insets*, detailed interaction networks of the individual subunits of AP-3, AP-4 and AP-5 (left) and of AP-1, AP-2 and TPC (right).

Next, we built the interactome network of AP-3 and AP-4 by means of their large subunits AP3B and AP4E and the small AP-4 subunit (AP4S) as baits for single-step AP-MS. The resulting AP-3 and AP-4 network consisted of 77 interactions among 54 proteins, demonstrating that, in addition to the interactions between core subunits within each complexes, AP-3, AP-4 and AP-5 were linked with each other (Fig. 2), because AP4E, AP4M and AP5M were identified with AP3B as a bait (Fig. 2, left inset). Markedly, P34 was also detected with both AP-4 subunits, in agreement with the identification of the AP4M subunit in the TAP-MS with P34 as a bait.

### Identification of the Arabidopsis AP-5 complex

Although sequence homology searches have identified three putative genes encoding AP-5 subunits in Arabidopsis, *AP5B* (AT3G19870), *AP5Z* (AT3G15160), and *AP5M* (AT2G20790)^2^, it is still unclear whether they function together as a *bona fide* complex in plant cells. To discover the core complex and the accessory proteins of AP-5, we used AP5B and AP5M as baits for single-step AP-MS. In total, 102 interactions among 59 proteins were present in the AP-5 network (Fig. 1 and Supplemental Data 1) and all three AP-5 subunits were identified together with the thus far uncharacterized proteins in plants ZINC FINGER FYVE DOMAIN-CONTAINING PROTEIN (ZFYVE) (AT2G25730) and SPATACSIN CARBOXY-TERMINUS PROTEIN (AT4G39420). Further sequence alignments revealed the homology of the SPATACSIN CARBOXY-TERMINUS PROTEIN and the ZFYVE protein to the mammalian SPG11 (with 60% similarity and 22.83% identity) and SPG15 (with 49% similarity and 22.57 identity) proteins, respectively, the two known accessory proteins of the AP-5 complex in mammalian cells^2, 49^. Consistent with the AP-3 and AP-4 interactome analyses, AP5M co-purified with four AP-4 subunits (Fig. 2, left inset). Taken together, these findings suggest the existence of crosstalk of AP-5 with AP-3 and AP-4 vesicle trafficking pathways next to the sub-clustering of the AP-1, AP-2 and TPC. For instance, the accessory adaptor protein SH3P2 was detected using AP3B, AP5B and CAP1 as baits. Interestingly, SH3P2 seems to function together in one complex with its homolog SH3P1, as shown through TAP-MS with SH3P1 as bait.

### Functional diversity of the AP complexes in Arabidopsis

To gain insight into the function of the AP complex-interacting partners, we analysed the Gene Ontology (GO) enrichment. All identified proteins were enriched in GO terms linked to protein transport and molecule metabolic process (Fig. 3), including “vesicle-mediated transport” and “establishment of protein localization”. The GO analysis of an individual AP complex revealed that the proteins associated with AP-1, AP-2, AP-3 and TPC were present in the “endocytosis”, whereas those of AP-4 and AP-5 were enriched in “lysosomal transport’’. The proteins concomitant with the accessory proteins P34, BAG4, AAK1, ECA1, ECA4 and CAP1 were paired with the terms “CME” and “clathrin coat assembly”. Additionally, the GO term “Golgi to vacuole transport” was shared by the AP-1, AP-4 and AP-5 interactors and the term “vacuolar transport” was associated with AP-3. Interestingly, specific AP complexes were coupled with some GO terms without direct links to vesicle trafficking. For example, some AP-1 interactors were connected with the terms “photoperiodism” and “mRNA export”. AP-2 and TPC shared the GO terms related to metabolic processes, whereas the term “response to metal ion” occurred only in AP-2. Specifically, AP-3 interactors were enriched in “cytokinesis” and “ribosome biogenesis”. By contrast, a cluster of terms related to “translation”, “protein complex disassembly” and “xylem development” were associated with AP-4 interactome. Proteins co-purifying with AP-5 were enriched, on the one side, in “lysosomal transport” and “Golgi to vacuole transport” and, on the other side, in “chromosome organization” and “histone modifications”. Altogether, the GO analysis revealed putative roles of AP complexes in different physiological pathways, seemingly not connected to endomembrane functions.

**Fig. 3.**
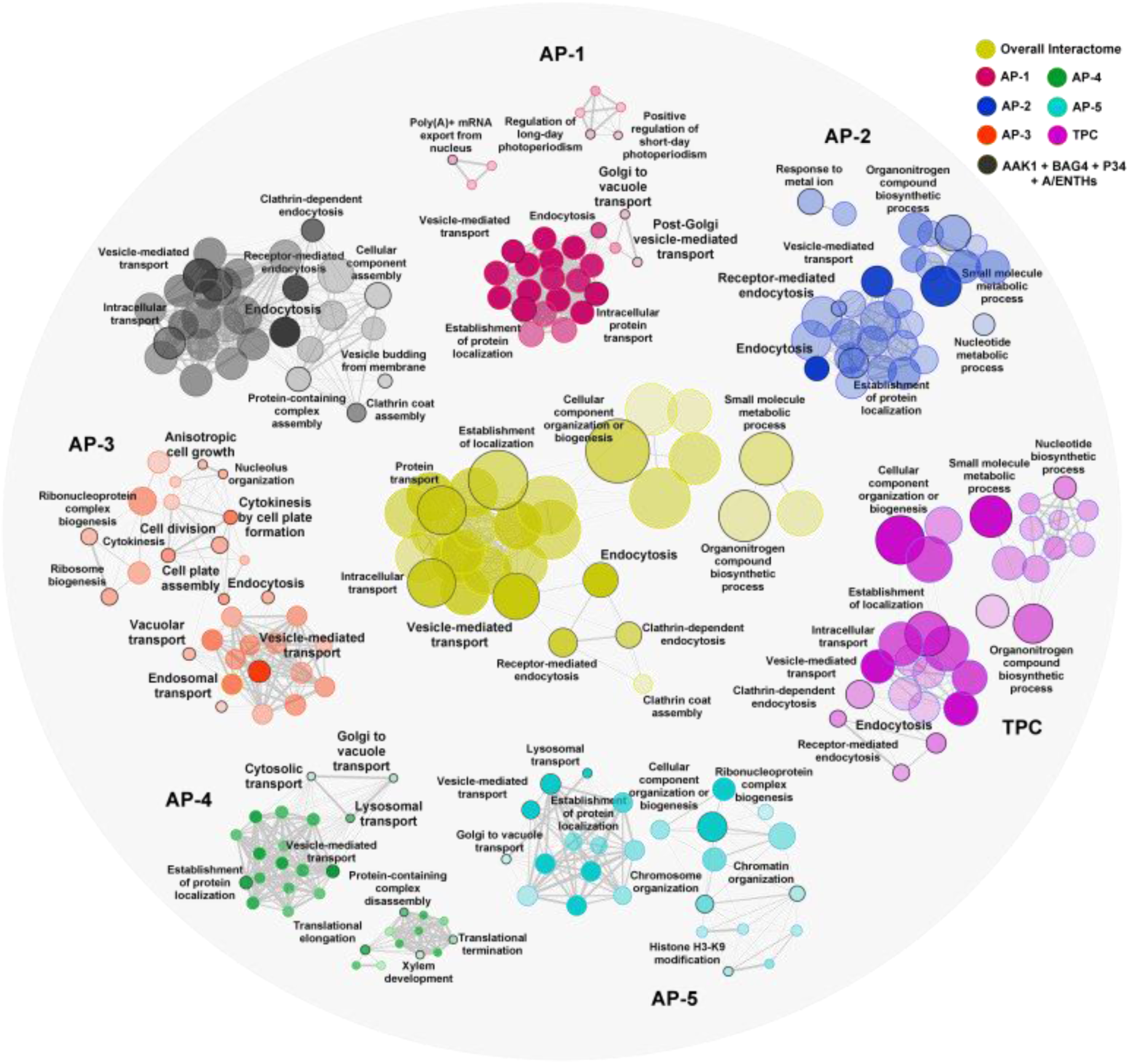
GO term enrichment analysis of the AP-associated proteins. Hierarchical clustering tree3 show the enriched gene sets associated with different AP complexes and accessory proteins. Each node indicates a significantly enriched GO term (FDR<0.05). The size and colour of the node represent the number of genes in the gene set and different complexes, respectively. The colour hue corresponds to the proportion of each cluster associated with the term. Two terms (nodes) are connected, when they share 20% or more genes. The line thickness indicates the number of shared genes between two terms. Selected GO terms (black outlines) are annotated.

### Validation of the AP interactomes

Since AP-5 had not been characterized in plants, we validated the interactions between the AP-5 subunits with co-immunoprecipitation (Co-IP) and ratiometric bimolecular fluorescence complementation (rBiFC) assays (Fig. 4a-c and Extended Data Fig. 4a). As predicted, AP5B and AP5M reciprocally co-immunoprecipitated when transiently co-expressed in tobacco (*Nicotiana benthamiana*) leaves (Fig. 4a). Similarly, AP5M co-immunoprecipitated with SPG11 and SPG15, whereas AP5B co-immunoprecipitated with SPG15 (Fig. 4a). In the rBiFC assay, co-expression of AP5M-nYFP (nYFP, the N-terminal fragments of YFP) with AP5B-cYFP (cYFP, the C-terminal fragments of YFP) and AP5Z-nYFP with AP5M-cYFP, but not with AP5B-cYFP, or nYFP-AP3B with AP5M-cYFP, resulted in fluorescence recovery in tobacco epidermal cells (Fig. 4b and Extended Data Fig. 4a). These data confirmed the interactions between AP-5 subunits and SPG proteins, supporting that AP-5 functions as a complex with SPG11 and SPG15 (Fig. 4c) in plant cells, similarly as in mammalian cells^46^.

**Fig. 4.**
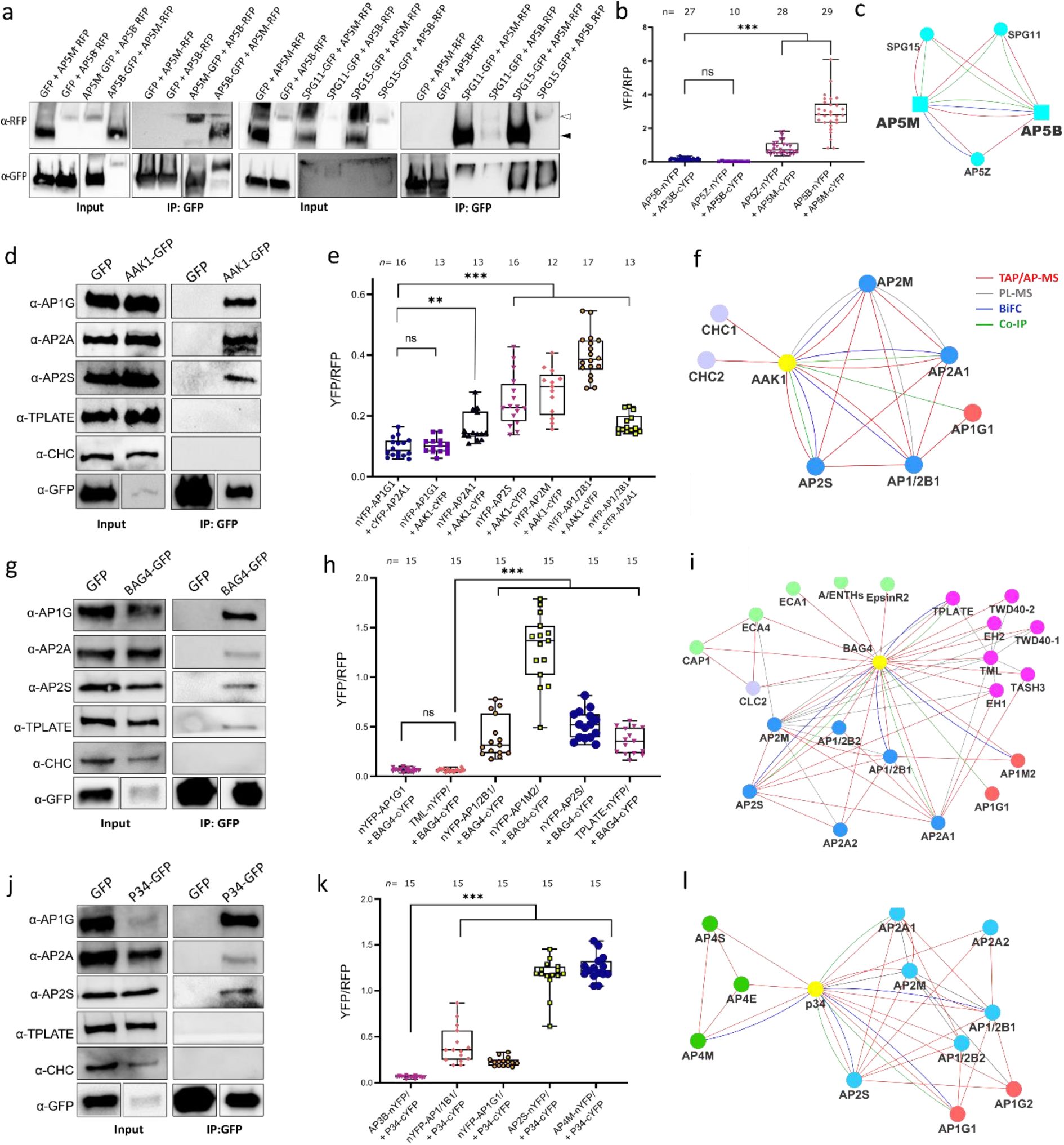
Validation of AP-5, AAK1, BAG4 and P34 protein interaction networks. **a**, Co-immunoprecipitation (Co-IP) assays in tobacco leaves transiently co-expressing different combinations of constructs. The *p35S:GFP* (GFP) construct was used as a negative control. White and black arrow heads indicate AP5B-RFP and AP5M-RFP, respectively. **b**, Quantification of rBiFC between AP5M, AP5B and AP5Z subunits. The interaction between AP5B and AP3B was used as a negative control. **c**, Cytoscape model of the AP-5 interactome. Edge colours represent the analysis methods used. **d, g, j**, Co-IP assays in Arabidopsis plants expressing *p35S:AAK1-GFP*, *p35S:BAG4-GFP* and *p35S:P34-GFP*. AAK1-GFP, BAG4-GFP and P34-GFP were pulled down with GFP-trap beads; AP-1, AP-2, TPC and clathrin were detected with α-AP1G, α-AP2A, α-AP2S, α-TPLATE and α-CHC antibodies. The *p35S:GFP* transgenic Arabidopsis plants were used as a negative control. Experiments were repeated 3 times independently with similar results. One representative experiment is presented. **e, h, k,** Quantification of the rBiFC (YFP/RFP ratio) between AAK1, BAG4, P34 and different AP subunits. AP1G1 and AP3B were used as negative controls for AAK1 and BAG4 and P34, respectively. The interaction between AP1/2B1 and AP2A1 was used as positive control. The rBiFC experiments were repeated twice of which one representative experiment is shown. *n,* number of cells analysed. \*\**P* ≤ 0.01; \*\*\**P* ≤ 0.001 (Brown-Forsythe and Welch ANOVA tests combined with Dunnett T3 multiple comparisons test), ns, not significant. **f, i, l,** Cytoscape models of AP-5, AAK1, BAG4 and P34 protein interaction networks. Edge colours represent the analysis methods used. Node colours represent the different complexes and protein families: AP-1, red; AP-2, blue; AP-4, green; clathrin proteins, magenta; A/ENTH proteins, light green.

Interactions between the accessory proteins AAK1, BAG4 and P34 and the AP subunits were also validated by Co-IP with transgenic Arabidopsis plants expressing *p35S:AAK1-GFP*, *p35S:BAG4-GFP*, *p35S:P34-GFP* and *p35S:GFP* constructs and by the rBiFC assay in tobacco leaf epidermal cells (Fig. 4d-l, Extended Data Fig. 4b, c, and Extended Data Fig. 5). AAK1 co-immunoprecipitated with subunits of AP-1and AP-2 but not with TPLATE and CHC (Fig. 4d). In the rBiFC assay, co-expressions of AAK1-cYFP with AP2A1-nYFP, nYFP-AP2M, nYFP-AP2S or nYFP-AP1/2B1 resulted in fluorescence signals (Fig. 4e and Extended Data Fig. 4b). Hence, AAK1 probably interacts directly with AP-2 (Fig. 4f) and the recovery of the AP1G subunit in the Co-IP of AAK1-GFP might result from the interaction with the shared AP1/2B1 subunit. Subsequently, Co-IP experiments in Arabidopsis seedlings revealed that BAG4 co-purifies with subunits of AP-1, AP-2 and TPC (Fig. 4g). These interactions were also confirmed by rBiFC (Fig. 4h and Extended Data Fig. 4c). Altogether, our data identified BAG4 as a hub protein associated with AP-1, AP-2 and TPC (Fig. 4i). In addition, P34 co-purified in Arabidopsis seedlings with subunits of AP-1, AP-2 and AP-4 (Fig. 4j and Extended Data Fig 5f), and rBiFC assays further corroborated the interaction between P34 and subunits of AP-1, AP-2 and AP-4 (Fig. 4k and Extended Data Fig. 5a). Interestingly, in the rBiFC assays, P34 also interacted with AP3D and AP5Z subunits (Extended Data Fig. 5b, c), suggesting that P34 binds to various AP complexes in rBiFC (Fig. 4l). However, P34 did not associate with AP3B via Co-IP in tobacco (Extended Data Fig. 5f). Furthermore, interactions between P34, BAG4 and AAK1 were also observed in rBiFC (Extended Data Fig. 5d,e). Interestingly, whereas the Arabidopsis NECAP-1 was identified in TAP-MS experiments with AP-1, rBiFC analysis revealed that it interacted with subunits of both the AP-1 and AP-2 complexes (Extended Data Fig. 6), similarly to the NECAP-1 in mammals^40^. The PL analysis with AP2M discovered several unknown PM-associated putative endocytic cargos and rBiFC (Extended Data Fig. 7) confirmed the interactions between AP2M and the cytokinesis-specific syntaxin KNOLLE^47^ and the FORMIN-LIKE6 (FH6) protein^48^

Taken together, we generated a highly reliable AP interaction network, including the composition of the AP-5, several AP complex accessory proteins, and revealed a number of regulatory proteins and putative cargoes associated with various AP complexes.

### The hub protein P34 regulates the stability of AP-1, AP-2 and AP-4

As the AP network had identified P34 as a hub protein interacting with several AP complexes, we investigated its function in Arabidopsis. Homozygous knockout *p34* mutants generated with the CRISPR/Cas9 technology (Extended Data Fig. 8a) could not be detected in the T1 generation nor in the offspring of two independent heterozygous lines, *p34-1(+/-)* and *p34-2(+/-)*. When grown in soil for 4 weeks, the rosette area of the heterozygous *p34-1(+/-)* and *p34-2(+/-)* mutants did not differ from the wild type (Extended Data Fig. 9a, b), but the siliques of the mature plants contained on average 25% abnormal seeds per silique, whereas only 0.5% of the seeds in the wild type siliques were abnormal (Extended Data Fig. 9c, d). The absence of homozygous *p34* mutant plants and the observed 25% seed abortion suggest that the *P34* gene is essential for embryogenesis. Screening the CRISPR/Cas9 population revealed one biallelic *p34* mutant, designed *p34-3(Δ/-)*. The *p34-3(Δ/-)* mutant contained one allele with a nucleotide inserted 63 bp after the start codon, leading to a stop after the 30^th^ amino acid and a second allele containing a 538 bp deletion and a 23 bp insertion, resulting in an in-frame translation of P34 with a deletion of 100 amino acids, from the 19^th^ to the 118^th^ amino acid (indicated as Δ) (Extended Data Fig. 8a, b). In contrast to the wild type and the two heterozygous *p34-1(+/-)* and *p34-2(+/-)* mutants, the *p34-3(Δ/Δ)* and *p34-3(Δ/-)* plants had smaller rosette areas when grown in soil (Extended Data Fig. 9a, b). The *p34-3(Δ/-)* mutant exhibited seed abortion rates similar to those of the heterozygous *p34-1(+/-)* and *p34-2(+/-)* alleles, whereas the *p34-3(Δ/Δ)* mutant behaved as the wild type (Extended Data Fig. 9c, d), suggesting that *p34-3(-)* and *p34-3(Δ)* <u>are</u> full knock-out and hypomorphic alleles, respectively. Examination of the primary root length of seedlings grown *in vitro* for 7 days revelled that it was significantly reduced for the *p34-3(Δ/-)* mutant only (Extended Data Fig. 9e, f). Further complementation analysis of the *p34-1(+/-)* and *p34-3(Δ/-)* mutants with the *pP34:gP34:GFP* construct were carried out and transgenic plants were recovered that resembled the wild type plants in terms of rosettes size, seed set and primary root length with the genotypes *pP34:gP34-GFP/p34-1(-/-)* and *pP34:gP34-GFP/p34-3(Δ/-)* (Extended Data Fig. 9a-f). As no homozygous *p34* mutant had been obtained and the biallelic *p34-3(Δ/-)* mutant did not allow genetic crossing, we generated β-estradiol-inducible *ipRPS5A>>CAS9-tagRFP-P34* knockout transgenic Arabidopsis plants (Fig. 5). Inference of CRISPR Edits (ICE) analysis of two independent T2 lines, *ipRPS5A>>CAS9-tagRFP-P34 #1-16* and *ipRPS5A>>CAS9-tagRFP-P34 #5-12*, hereafter designated *p34^iCRISPR^-1* and *p34^iCRISPR^-2*, respectively, showed high genome editing on *P34* (Extended Data Fig. 10a). In addition, the P34-GFP signal in the *p35S:gP34-GFP*/*p34^iCRISPR^-1* was almost eliminated after 5 days of β-estradiol treatment (Extended Data Fig.10b). When grown on agar medium containing 10 μM β-estradiol for 9 days, the primary root length of *p34^iCRISPR^-1* and *p34^iCRISPR^-2* plants were shorter than those of the DMSO controls and of the β-estradiol-treated wild type (Fig. 5a, b). Strikingly, the *p34^iCRISPR^* mutants also exhibited an agravitropic root growth after induction (Fig. 5c).

**Fig. 5.**
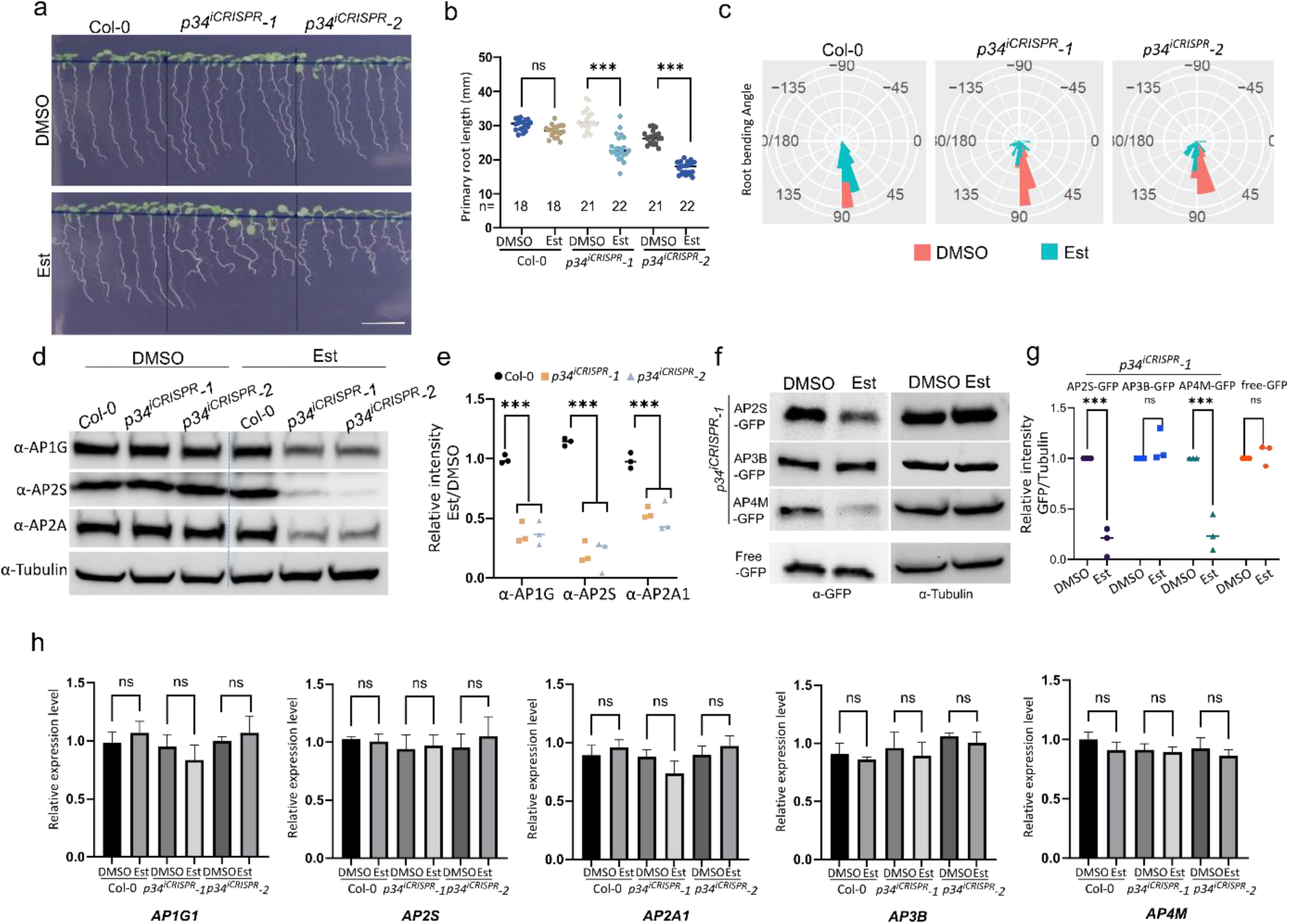
P34 regulates the stability of AP-1, AP-2 and AP-4. **a**, Wild type (Col-0), *p34^iCRISPR^-1* and *p34^iCRISPR^-2* seedlings grown on media with DMSO or 10 μM β-estradiol (Est) for 9 days. Scale bar, 10 mm. **b**, Primary root length of seedlings shown in (a). *n,* number of roots analysed. **c**, Root gravitropic bending of Col-0, *p34^iCRISPR^-1* and *p34^iCRISPR^-2* seedlings. Seven-day-old seedling grown on medium containing DMSO or 10 μM β-estradiol were gravistimulated by rotating 90° in two independent experiments (*n*=61-74 roots). **d, e**, Protein levels of AP2S, AP2A1, AP1G and tubulin of seedlings in (a) analysed with immunoblotting. Relative quantities of target proteins were normalized to tubulin for each sample. The values were further normalised to the DMSO control for each genotype. The experiments were repeated three times independently. One representative experiment is shown. The quantification in (e) combines three independent experiments. **f, g**, Protein levels of AP2S-GFP, AP3B-GFP and AP4M-GFP in *p34^iCRISPR^-1* background analysed in 9-day-old Arabidopsis seedlings treated with DMSO or β-estradiol. Seedlings expressing *35Sp:GFP* were used as control. Relative protein levels of AP2S-GFP, AP3B-GFP and AP4M-GFP were normalised to tubulin for each sample. The quantification in (g) combines three independent experiments. **h**, Expression levels of *AP1G1*, *AP2S*, *AP2A1*, *AP3B* and *AP4M* genes in 9-day-old seedlings. RT-qPCR analysis were done in three independent experiments each with three technical replicas. Transcript levels were normalised to *ACTIN2*. \*\*\**P* ≤ 0.001 [one-way ANOVA test in (b and h) and two-way ANOVA test in (e and g)], ns, not significant.

Recently, the human P34/AAGAB has been reported to control the assembly and stability of AP-2^42^. To establish whether this is the case in plants, we examined the protein levels of AP-1 and AP-2 in the inducible *p34^iCRISPR^-1* and *p34^iCRISPR^-2* mutants by immunoblots with α-AP1G, α-AP2A and α-AP2S antibodies (Fig. 5d-e). As anticipated, the AP1G, AP2A and AP2S protein levels were dramatically reduced in *p34^iCRISPR^* mutants. Similar results were obtained in the biallelic *p34-3(Δ/-)* mutant but the protein levels of CHC and TPLATE in *p34-3(Δ/-)* were similar to those of the wild type (Extended Data Fig. 10c-d). We further tested whether the protein levels of AP-3 and AP-4 were also affected in the *p34* mutants. To this end, we crossed the *p34^iCRISPR^-1* mutant with *pAP2S:AP2S-GFP/ap2s*, *pAP3B:AP3B-GFP/pat2-1*, and *pAP4M:AP4M-GFP/ap4m* and used the *p35S-GFP*/Col-0 plants as control. F2 seeds carrying the CRISRP/Cas9 construct were selected by seed fluorescence^49^ and used for further experiments. Immunoblot analysis indicated that the levels of AP2S-GFP and AP4M-GFP, but not of AP3B-GFP and GFP, were reduced after β-estradiol induction (Fig. 5f-g). No significant difference in the transcript levels of *AP1G1*, *AP2S*, *AP2A1*, *AP3B* and *AP4M* between the β-estradiol- and mock-treated plants were detected by RT-qPCR analysis (Fig. 5h). Taken together, our data show that P34 is an essential protein that controls the stability and possibly the assembly of AP-1, AP-2 and AP-4 but not of AP-3.

### P34 modulates various vesicle trafficking pathways

We next examined whether P34 modulates the trafficking pathways mediated by AP-1, AP-2 and AP-4 as their abundance was affected by the impaired P34 function. P34-GFP mainly localized to the cytoplasm in the root cells and was not visibly associated with membranes or organelles (Extended Data Fig. 10e). First, we followed the general endocytosis by the FM4-64 uptake in *p34^iCRISPR^-1* and *p34^iCRISPR^-2* seedlings after β-estradiol induction and in the *p34-3* mutants. The *ap2m-2* mutant was used as a reference. As expected, the internalization of FM4-64 in the root cells was significantly reduced in the *p34^iCRISPR^* mutants (Fig. 6a, b). In addition, *p34-3(Δ/-)* also showed a decreased internalization of FM4-64 when compared to the wild type (Extended Data Fig. 10f-g).

**Fig. 6.**
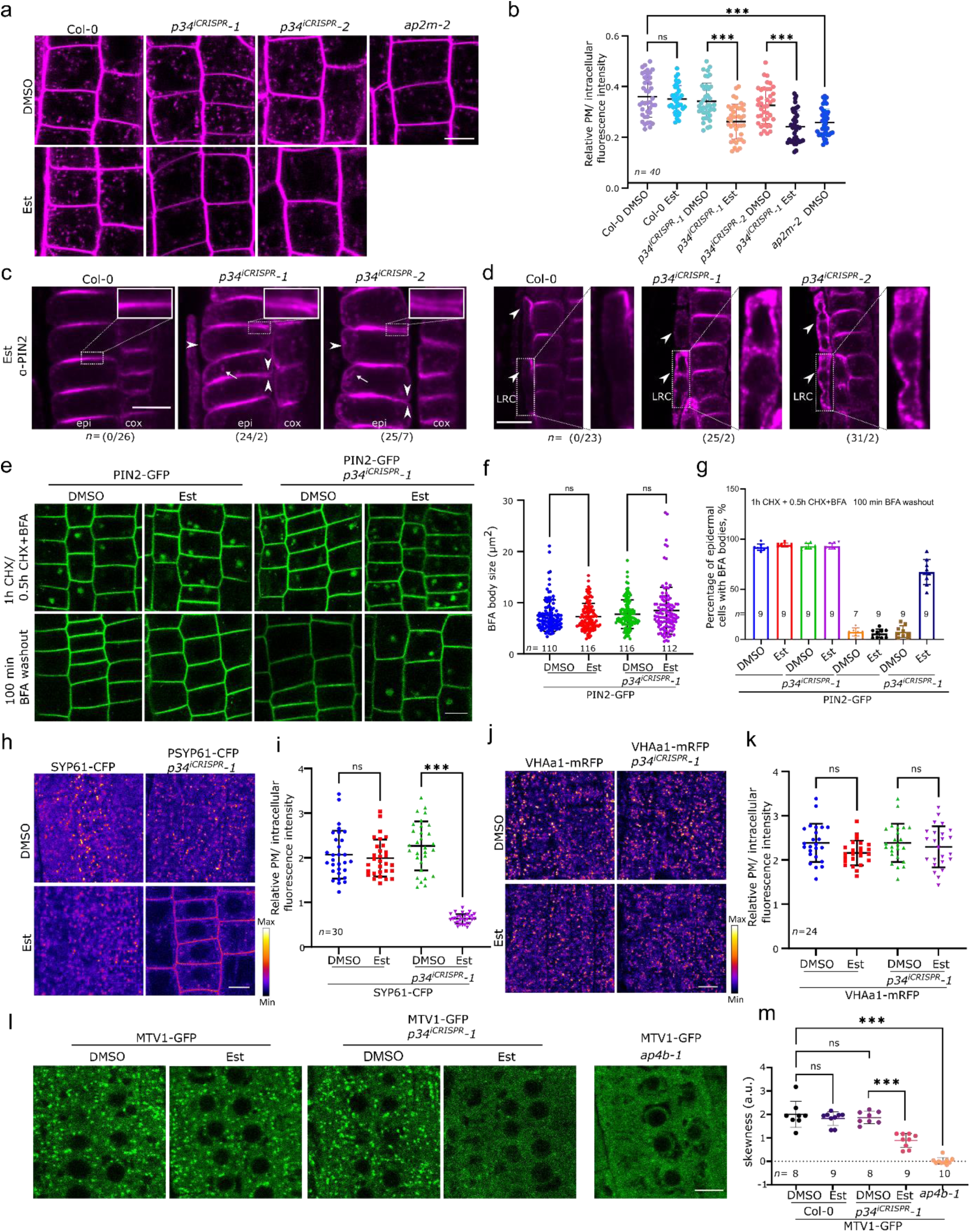
P34 regulates vesicle trafficking depending on AP-1, AP2 and AP-4. **a**, FM4-64 uptake in the *p34^iCRISPR^-1* and *p34^iCRISPR^-2* mutants. Root epidermal cells of 5-day-old wild type (Col-0), *p34^iCRISPR^-1*, *p34^iCRISPR^-2* and *ap2m-2* seedlings grown on DMSO or 10 μM β-estradiol (Est) were imaged after staining with FM4-64 (2 μM, 15 min). **b,** Relative intracellular to plasma membrane (PM) fluorescence intensity ratio of FM4-64 of images in (a). *n*=40 cells in 10 roots were analysed. **c, d,** Immunolocalization of PIN2 in epidermal (epi) and cortex (cox) cell in (c) and in lateral root cap (LRC) cells in (d) of 7-day-old Col-0, *p34^iCRISPR^-1* and *p34^iCRISPR^-2* roots treated with 10 μM β-estradiol. Arrow heads and arrows indicate the non-polar distribution of PIN2 in epidermal and lateral root cap cells and endosomal PIN2, respectively. The insets show the non-polar distribution of PIN2 in the epidermis. Scale bars, 10 µm. The ratio of PIN2 abnormal/normal localization in roots is indicated for three independent experiments. PIN2-GFP in the root cells of 5-day-old wild type and *p34^iCRISPR^-1* seedlings grown on DMSO and 10 μM β-estradiol. The seedlings were pretreated with cycloheximide (CHX) (50 μM, 1 h) and then treated with CHX (50 μM) and Brefeldin A (BFA) (50 μM) for 30 min. Seedlings were washed with media containing CHX (50 μM) and imaged 100 min after the wash. **f**, BFA body sizes before the washout. *n,* number of BFA bodies. At least seven roots were imaged and measured. **g**, Percentage of BFA body-containing cells in PIN2-GFP seedlings before and after BFA washout in (e). *n*, number of roots analysed. **h**, Fluorescence intensity images of SYP61-CFP in the root cells of 5-day-old Col-0 and *p34^iCRISPR^-1* grown on DMSO or 10 μM β-estradiol. **i**, Relative intracellular to PM fluorescence intensity ratios of SYP61-CFP in one representative experiment. ns, not significant. *n,* number of cells being analysed. At least five roots were imaged and measured. **j**, Fluorescence intensity images of VHAa1-mRFP in the root cells of 5-day-old Col-0 and *p34^iCRISPR^-1* grown on DMSO or 10 μM β-estradiol. **k**, Relative intracellular to PM fluorescence intensity ratios of VHAa1-mRFP in one representative experiment. *n,* n<u>u</u>mber of cells analysed. **i,** Localization of MTV1-GFP in the root cells of 5-day-old Col-0, *p34^iCRISPR^-1* and *ap4b-1* grown on DMSO or 10 μM β-estradiol. **m**, Quantification of the presence of defined MTV1-GFP punctate signals by analysis of the skewness of the signal intensity from at least seven independent point populations (seven roots). Higher and lower skewness indicate and more diffuse signals, respectively. The imaging of all genotypes was repeated three times. One representative experiment is shown. Error bar represents standard deviation. \*\*\**P* ≤ 0.001 [one-way ANOVA in (b, f, i, k, m)], ns, not significant, a.u., arbitrary units.

Subsequently, the localization of the known CME cargo PIN2^11, 37^ in root cells of the *p34^iCRISPR^* mutants was studied by whole-mount immunofluorescence labelling (Fig. 6c, d). The polar localization of PIN2 was severely disrupted, whereas its endosomal localization was more pronounced in the epidermal and lateral root cap cells. We also examined the accumulation of PIN2-GFP in Brefeldin A (BFA)-induced endosomal aggregates (so-called BFA bodies). No significant differences in PIN2-GFP BFA body size was observed between the wild type and the *p34^iCRISPR^* mutants after a 1-h BFA treatment (50 µM) (Fig. 6e, f). In contrast, the dissipation of BFA bodies was significantly slower that in the *p34^icrispr^* mutant than in the wild type, as indicated by the retention of abundant BFA bodies in the *p34^icrispr^* mutant after a 100-min BFA washout (Fig. 6e, g). These data indicate that P34 is essential for endocytosis, recycling, BFA-sensitive endosomal aggregation and polarity establishment of PIN2, probably through regulation of the AP-2 and AP-1 stability in the cytoplasm.

As AP-1 affects exocytosis, endocytosis and vacuolar trafficking^50^, we investigated whether impairing the P34 function influences the trafficking of the secretory marker, secRFP^51^, the PIN2^37^ and KNOLLE^7, 47^ proteins, and the localization of the TGN/EE-resident proteins, SYNTAXIN 61 (SYP61)^56^ and VACUOLAR ATPASE SUBUNIT a1 (VHAa1)^50^. To this aim, we assessed the secretion of secRFP to the apoplast in *p34^iCRISPR^-1*. The localization of secRFP in the *p34* mutants after β-estradiol induction remained apoplastic (Extended Data Fig. 11a), indicating that the secRFP secretion to the apoplast was not affected. By means of whole-mount immunofluorescence labelling, we determined the localisation of KNOLLE during cytokinesis. In the wild type, KNOLLE is recruited to the developing cell plate via the secretory pathway^7^, is targeted for vacuolar degradation once cytokinesis is completed^52^ and is retrieved from the cell plate insertion site by endocytosis^25, 53^. Although in the *p34^iCRISPR^-1* plants some KNOLLE reached the cell plate during and post-cytokinesis, much more KNOLLE was detected in endosomes, possibly reflecting partial defects in its secretion and/or vacuolar trafficking (Extended Fig. 11b). Visualization of PIN2-GFP signals after dark treatment revealed that the PIN2-GFP accumulation in the vacuole was strongly reduced in the β-estradiol-treated *p34* mutant similarly to the *ap1m2-1* mutant^7^ (Extended Data Fig. 11c, d), further indicating that besides secretion, vacuolar trafficking was also affected in the *p34*. Furthermore, when grown on DMSO, SYP61-CFP mainly localized to the TGN/EE in both wild type and *p34^iCRISPR^-1* root cells (Fig. 6h-i). However, after β-estradiol induction SYP61-CFP was mislocalized to the PM in *p34^iCRISPR^-1* line (Fig. 6h-i), likewise as when SYP61 was examined in the *ap1m2* mutant^50^. Moreover, the localization of VHAa1-RFP in *p34^iCRISPR^-1* after β-estradiol induction was not significantly altered (Fig. 6j-k) thus resembling the VHAa1 localization in the *ap1m2* mutant^50^. Taken together, these results suggest that in the *p34^iCRISPR^* mutants some but not all, aspects of AP-1-mediated trafficking are affected.

Since impairment of the P34 function had an impact on the AP-4 stability, we studied the localization of the Epsin-like protein MODIFIED TRANSPORT TO THE VACUOLE1 (MTV1), previously shown to localize to the TGN/EE in AP-4-dependent manner^23^, in the *p34^iCRISPR^-1* line (Fig. 6m, n). Similarly to its expression in the *ap4b-1* mutant, the punctate pattern localization of MTV1-GFP in the *p34^iCRISPR^-1* mutant was reduced after β-estradiol induction (Fig. 6l-6m). Notably, in contrast to that in the *ap3* mutant^18^, the vacuolar morphology remained unaffected and PIN2-GFP did not accumulate abnormally in the vacuole in *p34^iCRISPR^-1* under normal light conditions (Fig. 6c, Extended Data Fig. 11e), suggesting that the AP-3-depnedent pathway is functional.

In summary, our data demonstrate that P34 regulates CME to some extent, but had a more pronounced impact on the trafficking to the vacuole, the localization of a subset of TGN/EE proteins, such as SYP61 and the AP-4 cargo MTV1, but not on the default secretion to the apoplast. We hypothesise that these result are achieved through maintenance of the correct protein levels of AP-1, AP-2 and AP-4 and possibly of their complex assembly.

## Discussion

AP complexes are key regulators of the intracellular protein transport. AP dysfunction in mammalian cells disturbs many cellular processes, including organelle dynamics, tissue homeostasis and signal transductions, and provokes various heritable diseases and neurological disorders^3^. In plants, AP complexes also play critical roles in protein sorting and regulate several physiological processes^54^. Despite many similarities between plants and mammals, some structural differences make plant vesicle trafficking fundamentally distinct^4^. Firstly, the secretory, but the endocytic pathway as well, converge at the plant TGN/EE. Secondly, although plant cells rarely contain lysosomes, they possess large vacuoles with multiple functions. Thirdly, supposedly aiding to overcome the high-turgor pressures, an actin-independent endocytic pathway combined with an additional membrane bending machinery, such as TPC, is evolutionarily retained in plants^4, 26^. These differences imply plant-specific adaptations to control the vesicle trafficking mechanisms.

Mammalian AP-1, AP-2 and AP-3 interact with clathrin, whereas AP-4 and AP-5 do not^3^. However, only AP-1 and AP-2 are enriched in the mammalian CCVs and participate in their formation^3^. Although AP-3 can interact with clathrin, this interaction is currently viewed as non-essential under physiological conditions^55^. AP-5 does not dependent on clathrin and, instead associates with SPG11 and SPG15, which have been predicted to have α-solenoid structures similar to those of CHC, TPC and coatomer protein complex (COPI) subunits^56^. On the contrary, all subunits of AP-1, AP2 and AP-4, but not those of AP-3 or AP-5, are enriched in plant CCVs^35^. The PL with TML identified CLC2 that, in agreement with previous reports, connects TPC with clathrin^25^. In our interactome, however, no clathrin associated with any of the five AP complexes. Conversely, the TAP-MS detected CHC1 and CHC2 associated with the AP-2 binding partner, AAK1, whereas CLC1 and CLC2 associated with the AP-1 and AP-2 binding partner, BAG4. Nevertheless, these interactions were not confirmed by the Co-IP experiments.

Human AAK1 regulates CME by phosphorylating the medium subunit of AP-2, which promotes binding of AP-2 to cargo proteins during CCV formation^44, 57^. This kinase modulates several pathways, including NOTCH and WNT signalling^44, 45, 58^. The AP-2 network showed that the Arabidopsis AAK1 binds specifically to AP-2, indicating that AAK1 might regulate AP-2 functions and CME in plant cells. AAK1 also interacted with BAG4, which had been identified as a hub protein associated with AP-1, AP-2 and TPC. The BAG family is evolutionarily conserved in eukaryotes, and the human BAG proteins act as co-chaperones of the heat shock protein (HSP70) family. BAG4 proteins are involved in multiple functions, such as apoptosis, autophagy, proteasomal and lysosomal degradation, neurogenesis and tumorigenesis^59^. Arabidopsis contains seven BAG family members and BAG4 was shown to be involved in salt responses and stomatal movement^41^. Consistent with the obtained protein-protein interaction data, BAG4 also associated with clathrin, TOM1-LIKE (TOL) ESCRT-0 proteins^60, 61^, two ANTH-type clathrin adaptors, ECA4 and CAP1^25^ and a group of heat shock proteins^41^, suggesting a role for the Arabidopsis BAG4 in protein degradation either linked to autophagy, the ubiquitin-mediated proteasomal degradation pathway or endocytosis^61, 62^. Taken together, our data uncover an intimate network of the AP-1, AP-2, TPC, AAK1, BAG4 and the A/ENTH proteins that are involved in clathrin-dependent protein trafficking.

The AP interactome revealed that in Arabidopsis AP-3, AP-4 and AP-5 are more closely related to each other, not only because they share several interacting components but also because some of their subunits co-purified. Instead of acting in a clathrin-dependent manner, the human AP-5 cooperates with SPG11 and SPG15 as a scaffold and a docking site, respectively^46^. We discovered that Arabidopsis AP-5 also associates with the plant homologues of the human SPG11 and SPG15, AT2G25730 and AT4G39420, respectively, of which the functions in plants are still not elucidated. In spite of the high probability that the Arabidopsis AP-5 also acts together with SPG11 and SPG15 in plants, we could not exclude that AP-5 might utilize or share subunits with AP-3 and/or AP-4, because the AP5S subunit is absent and the three AP complexes crosstalk during the trafficking between TGN and late endosomes in plant cells.

Furthermore, we discovered one important regulatory protein that links the functions of AP-1, AP-2 and AP-4 in plant, the Arabidopsis P34. In humans, P34/AAGAB controls the sequential assembly of AP-1 and AP-2, but not that of AP-3^42^, and it is still unclear whether P34 regulates AP-4 or AP-5. The Arabidopsis P34 positively regulated the stability of AP-1 and AP-2, whereas the AP-3 stability remained unchanged in the *p34* knockout mutants, consistently with the results in human cells. In Arabidopsis, P34 also interacted with the AP-4 subunits (AP4E, AP4M and AP4S) and the AP4M protein was significantly reduced in the *p34* mutant. In contrast, P34 did not co-purify with TPC, implying that in plants it directly regulates the protein stability of the three clathrin-associated AP complexes, AP-1, AP-2 and AP-4^35^ but not that of AP-3 and TPC. Therefore, P34 might be a regulator of multiple clathrin-mediated vesicle trafficking pathways in Arabidopsis. Although debatable, some evidence exists that the activities of clathrin-dependent post-Golgi trafficking and CME are highly coordinated in plant cells^50, 63^. Recently, the recruitment of several endocytic components to the PM, including CLC1, CHC1, AP2M, AP2S, TPLATE and TML, was shown to be reduced in the *ap1m2* mutant, but their protein levels to remain unchanged^50^. Similarly, in mutants defective in AP-2 and TPC-dependent CME, the clathrin and AP-1 recruitment to the TGN/EE as well as exocytosis are significantly impaired^50^. As the impaired P34 function strongly affected the protein levels of the AP-1, AP-2, AP-4 subunits and probably other CME components we hypothesized that multiple trafficking pathways would be affected in the *p34* mutant. Indeed, as predicted, the internalization of the lipophilic PM tracer dye FM4-64 was significantly delayed in the *p34* mutant and the localization of the TGN/EE-resident protein SYP61 was altered similarly to that in *ap1m2*^50^, suggesting that P34 regulates the trafficking pathways controlled by the AP-1 and AP-2. Why SYP61 is mistargeted to the PM when the AP-1 function is impaired remains to be determined, but in absence of degradation, SYP61might be assumed to be targeted by default to the PM. PIN2 polarization and recycling from BFA-sensitive endosomes was also defective in *p34*. Previously, modification of the PIN2 polar localization in the cells of staminal filaments of *ap2m-1*^11^, and decrease in the recycling of PIN2 from the BFA-sensitive endosome in mutants of AP-1 and AP-2 subunits have been shown^9, 50^. In addition, a more diffuse endosomal KNOLLE signal was observed in *p34,* reminiscent of KNOLLE localization in *ap1m2-1*^8^. The lack of ectopic KNOLLE localization at the post-cytokinetic cell wall insertion site indicates that *p34* is a weaker mutant than *ap1m2*, without impact on sterol- and TPC-dependent endocytosis^53, 64^. Consistently, the secretory marker secRFP was correctly transported to the apoplast in *p34*, but was impaired in the *ap1m2* mutants^7, 50^. Previously, AP-1 and AP-4 have been reported to localize in different subdomains of TGN/EE^23^, and to initiate secretory and vacuolar trafficking from different subregions of TGN/EE^5^. P34 directly regulated the function of AP-4, because the TGN/EE-resident protein MTV1 that requires AP-4 function for its loclaization^23^, was mislocalized and mostly remained cytosolic in *p34*. The differences between the *p34* and *ap1* mutants might therefore be linked to the TGN integrity that might be more affected in *ap1* than in *p34*.

In summary, the generated AP interactome provides useful resources to discover unknown regulators or co-operators of the AP complexes that might be unique to plant cells or have a common function in all eukaryotic cells, such as AAK1, P34, SPG11, and SPG15. Our findings identified P34 as a regulator of clathrin-mediated post-Golgi transport and CME by abundance maintenance of the AP-1, AP-2 and AP-4 complexes in plant cells. Which factors regulate the stability and possibly the assembly of TPC, AP-3 and AP-5 in Arabidopsis remains to be determined.

## Methods

### Plant material and growth conditions

*Arabidopsis thaliana* (L.) Heynh., accession Columbia-0, plants were used for all experiments. *Arabidopsis* seeds were sown on half-strength Murashige and Skoog (½MS) agar plates without sucrose or in liquid ½MS without sucrose and vernalized at 4°C under dark conditions for 3 days. Seeds were germinated and grown at 22°C and a 16-h light/8-h dark photoperiod for 5, 7 or 9 days, according to the experimental purposes. For *P34*-inducible CRISPR editing experiments, seeds were germinated and grown directly on ½MS containing either DMSO or 10 µM β-estradiol at 22°C and a 16-h light/8-h dark photoperiod for 5, 7 or 9 days. Transgenic *Arabidopsis* lines expressing *p35S:GFP/Col-*0^65^, *pAP2S:AP2S-GFP/ap2s*^15^, *pMTV1:MTV1-GFP/Col-0*^23^, *pMTV1:MTV1-GFP/ap4b-1*^23^, *pAP4M:AP4M-GFP/ap4m-2*^66^, *pAP3B:AP3B-GFP/pat2-1*^18^*, pSYP61:SYP61-CFP*^67^, *pVHAa1-VHAa1-mRFP*^68^, *p35S:secRFP*^51^, *pPIN2:PIN2-GFP*^37^, *ap2m-2*^16^ have been previously described. Wild-type tobacco (*Nicotiana benthamiana*) plants were grown in the greenhouse under a normal 14-h light/10-h dark regime at 25°C.

### Rosette area measurements

Four-week-old plants grown in soil in multipot trays were imaged. The total leaf area of individual plants was selected by means of the ImageJ colour threshold tool. Per genotype, 8-9 seedlings were planted randomly to exclude position effects. The experiments were repeated three times independently. The rosette area from a total of 24-25 plants per genotype was measured.

### Generation of constructs and transformation

For the entry clones, genomic DNA of *Arabidopsis AP1G1*, *AP1G2*, *AP1S1*, *AP2S*, *AP3B*, *AP4E*, *AP4S*, *AP5B*, *AP5M*, *P34*, *BAG4*, *AAK1*, *SH3P1*, *NECAP-1*, *ECA1*, *ECA4* and *CAP1* were cloned into a Gateway entry vector *pDONR221* with a BP reaction. For the TAP-MS and AP-MS constructs, the above genes were fused N-terminally or C-terminally to the GS^TEV86^ or GS^rhino87^ TAP tag under control of the *35S* promoter by subcloning the entry clones into *pKCTAP* or *pKNGSrhino/pKNGSTAP*^69^ with a Gateway LR reaction. For *p35S:AAK1-GFP*, *p35S:BAG4-GFP, p35S:P34-GFP* transgenic *Arabidopsis*, the genomic DNA of *AAK1* and *P34* and the cDNA of *BAG4* were first cloned into *pDONR221,* then subcloned into the binary vector *pK7FWG2*^70^ by a LR reaction and finally transformed into *Arabidopsis* Col-0 by floral dip. For the Co-IP experiments, entry clones of *AP3B* and *AP4E* were subcloned into *pK7FWG2*^70^ to generate the *p35S:AP3B-GFP* and *p35S:AP4E-GFP* constructs. For the *p35S:P34-mCherry* construct, cDNA of *P34* was cloned into *pGGB000* and subcloned into the backbone vector *pFASTRK-AG* by the Golden Gate system with *pGG-A-35SP-B, pGG-C-mCherry-D* and *pGG-D-35ST-G*^71^. To generate the *pP34:gP34-GFP* construct, the promoter (1812 bp) and genomic DNA (2035 bp) of *P34* were cloned into the *pDONR P4-P1r* and *pDONR221* vectors, respectively, and then subcloned into *pH7m34GW* vector with *pDONR P2R-P3-EGFP* by means of the MultiSite Gateway system. The rescued lines were generated by transforming the *pP34:gP34-GFP* construct into *p34-1(-/+)* and *p34-3(Δ/-)* plants by floral dip.

For the CRISPR/Cas9 constructs, two gRNAs that targeted the first and the last exon of *P34* were designed with the ‘CCtop’ software (https://cctop.cos.uni-heidelberg.de:8043/)^72^. The two gRNAs (Supplementary Table 1) were cloned into the entry vectors *pGG-C-AtU6-26-BbsI-ccdB-BbsI-D* and *pGG-D-AtU6-26-BbsI-ccdB-BbsI-E* according to the previously described protocol^71^. The CRISPR/Cas9 expression construct was generated by assembling the entry clones with *pGG-A-linkerIII-C*, *pGG-E-linker-G*^92^, into the backbone vector *pFASTRK-Atcas9-AG*^49^ with the Golden Gate reaction. For the inducible CRISPR construct, the entry clones of the gRNAs and *pGG-A-linkerIII-C*, *pGG-E-linker-G* were first cloned into the *pEN-R2-AG-L3*^74^ with the Golden Gate reaction. The resulting *pEN-R2L3-gRNA* constructs were then subcloned with *pEN-R4-RPS5A-XVE-L1*^62^ and *p221z-CAS9p-tagRFP* into the *pK8m43GW* binary vector by a Multisite Gateway reaction. The CRISPR and inducible CRISPR expression constructs were then transformed into Col-0 by floral dip. For genotyping, the gnomic DNA sequence of *P34* of the transgenic plants were amplified by PCR with the primers listed in the Supplementary Table 1.

For the rBiFC assay^75^, cDNA of *BAG4, P34, AAK1, AP1M2, AP4M, AP5M, AP5Z, AP5B, FH6, KNOLLE* and *BIN2* were first cloned into *pDONR-p3p2* or *pDONR-p1p4* to generate the entry clones. The entry clones were then subcloned with *pDONR-p3p2-AP2S, pDONR-p3p2-AP1G1* or *pDONR-p3p2-AP2A1, pDONR-p3p2-AP12B1* and *pDONRp3p2-AP2M*^13^ into the expression vectors *pBiFC-2in1-NN, pBiFC-2in1-CN, pBiFC-2in1-NC or pBiFC-2in1-CC*^75^ by Multisite Gateway cloning. All primers used for cloning are listed in the Supplementary Table 1.

### Chemical Treatments

β-estradiol (Sigma-Aldrich, 20 Mm stock in DMSO), BFA (Sigma-Aldrich, 50 mM stock in DMSO), CHX (Merck, 50 mM stock in DMSO), and FM4-64 (Invitrogen, 2 Mm stock in water), were used at the concentrations indicated.

### rBiFC analysis and quantification

For rBiFC assay, the *Agrobacterium tumefaciens* strain C58 carrying the 2in1 constructs were coinfiltrated with a p19-carrying strain into *Nicotiana benthamiana* leaves and incubated in the greenhouse for 2 days. Afterwards, infiltrated leaves were observed either with an SP8 (Leica) or Fluoview1000 (Olympus) confocal microscope. The rBiFC analysis was quantified by means of 8-bit greyscale images containing fewer than 1% saturated pixels. The PM was outlined with the selection brush tool in Fiji and the mean intensity of the selected region of interest was calculated in both channels to generate an YFP/RFP ratio per cell as previously descibed^75^. All rBiFC combinations tested are listed in Supplementary Data 2.

### Tandem affinity purification, single-step affinity purification and proximity labelling experiments

All GS^TEV^ and GS^rhino69^ TAP tag fusion constructs were transferred to the *Agrobacterium tumefaciens* C58C1 strain pMP90 for transformation into the PSB-D *Arabidopsis* cell culture. The transformed *Arabidopsis* cells were cultured and harvested, followed by protein complex purification and LC-MS/MS identification of the purified proteins as previously described^69^. For proximity biotinylation experiments, AP2M and TML were combined C-terminally with TurboID and a 65-amino-acid linker sequence in between^36^. *Arabidopsis* cells were transformed according to the procedure for the TAP purifications. Biotinylated proteins were purified and identified as described previously^36^, by growing the cell cultures at 28°C, incubating them for 24 h with 50 µM exogenous biotin, extracting the proteins under very harsh conditions (100 mM Tris [pH 7.5], 2% [w/v] SDS and 8 M Urea) and a two-step elution, followed by LC-MS/MS analysis on a Q Exactive (Thermo Fisher Scientific).

### TAP-MS, AP-MS and PL-MS analysis and background filtering

Two and three replicates were done for TAP and single-step AP, respectively. After each set of two TAP or three AP purifications, the identified protein list was filtered versus a list of nonspecific proteins, assembled from a large dataset of TAP-MS or AP-MS experiments as previously described^69^. All proteins appearing with three or more bait groups in the large dataset were included in a list of nonspecific proteins. For each TAP-MS set of two experiments to be analysed, the background was marked in the list of the copurified proteins. To prevent that true interactors were filtered out because of their presence in the list of nonspecific proteins, the ratio was calculated of the average normalized spectral abundance factors (NSAFs) of the identified proteins in the bait pull-downs versus the corresponding average NSAFs in a large control set of pull-downs. Proteins identified with at least two peptides in at least two experiments, which did not occur in the background list, or showed a high (at least 20-fold) enrichment versus the large dataset of TAP experiments, were retained. Identifications without high enrichment versus the large dataset were removed, unless they were present in one of the other TAP-MS experiment sets. For each AP-MS set of three experiments to be analysed, the same approach was followed, additionally with a Student’s *t* test. Identifications with a high (NSAF ratio ≥ 10) and significant (-log(*P* value) ≥ 10) enrichment versus the large control AP-MS dataset were retained. The PL-MS data were filtered by comparison with a PL-MS control dataset built from experiments done under the same conditions. Identifications with enrichment score ≥ 20 were considered enriched, namely enrichment score (ES) = NSAF ratio x -log(*P* value). Protein interactions were visualized in networks with the Cytoscape v3.7.2 software using the input in Supplementary Dataset 1 or in a dot plot matrix with ProHits-viz^76^.

### Gene set enrichment analysis

Gene set enrichment was analysed with ShinyGO v0.741:Gene Ontology Enrichment Analysis + more (http://bioinformatics.sdstate.edu/go/)^77^ in *Arabidopsis* with a *P* value cut-off (false discovery ratio [FDR]) = 0.05 and the top 30 pathways. The networks of enriched GO terms were visualized with the Cytoscape v3.7.2 software. Two terms were connected when they shared 20% or more genes. The fold change of each term was indicated by the size of the nodes and the number of shared genes between two terms was indicated by the line thickness.

### Immunohistochemistry

Immunolocalizations were carried out on 7-day-old or 9-day-old seedlings grow on ½MS medium containing 10 µM β-estradiol by means of the immuno-robot InsituPro Vsi II (Intavis), as previously described^78^. In brief, the samples were fixed by paraformaldehyde (4% [v/v]) in phosphate-buffered saline (PBS) for 1 h in vacuum. After fixation, the samples were transferred to the robot and subsequently subjected to 6 washes (3 times in PBS, pH 7.4, and 0.1% [v/v] Triton X-100 [PBS-T] and 3 times in distilled water with 0.1% [v/v] Triton X-100; 5 min each), cell wall digestion (1.5% [w/v] driselase in PBS, for 30 min at 37°C), 3 washes (PBS-T, for 5 min), permeabilization (3% [v/v] NP-40, 10% [v/v] DMSO in PBS; for 30 min at room temperature), 3 washes (PBS-T, for 5 min), blocking solution (3% [w/v] bovine serum albumin in PBS, for 1 h at 37°C), primary antibody solution (primary antibodies diluted in blocking solution, for 4 h at 37°C), 5 washes (PBS-T, for 5 min), secondary antibody solution (secondary antibodies diluted in blocking solution, for 3 h at 37°C), 5 washes (PBS-T, for 5 min); when 4′,6-diamidino-2-phenylindole (DAPI) staining was needed, samples were incubated in 1 µl DAPI in stock with 5 ml distilled water for 15 min, followed by 5 washes with distilled water for 5 min). The dilutions of the primary antibodies were: rabbit α-PIN2 (1:600)^92^, mouse α-Tubulin (Sigma-Aldrich, T5168) (1:1000), rabbit α-Knolle (1:1000)^52^ (kind gift from Dr. Gerd Jürgens) and of the secondary antibodies: AlexaFluor488 goat α-mouse (1:600) (A-11001, Thermo Fisher Scientific) and AlexaFluor555 donkey α-rabbit (1:600) (A-31572, Thermo Fisher Scientific). The DAPI (D9542, Sigma-Aldrich) stock concentration was 1 mg/ml.

### Gravitropic responses

Five-day-old Col-0, *p34^iCRISPR^-1* and *p34^iCRISPR^-2* seedlings, light-grown vertically on medium supplemented with DMSO or 10 μM β-estradiol, were gravistimulated by a 90° rotation. The angle of the root tips deviating from the vertical plane was measured with the ImageJ (http://rsb.info.nih.gov/ij/) software 24 h after gravistimulation. Gravicurvature was quantified in two independent experiments with in total 61 to 74 roots. All gravitropically stimulated roots were assigned to a gravitropism diagram by an online tool (https://rmtrane.shinyapps.io/RootNav/).

### Quantitative RT-PCR

RNA was extracted from 9-day-old seedlings with the ReliaPrep™ RNA Miniprep Systems (Promega). Of purified RNA, 1 μg was amplified in a reverse transcriptase reaction with the qScript XLT 1-Step RT-PCR Kit (Quantabio). Subsequently, qPCR was run with the SYBR Green master mix (Roche) with gene-specific primers designed to amplify *AP1G1*, *AP2S*, *AP2A1*, *AP3B* and *AP4M*. *ACTIN2* was used as the normalization reference. Primers are listed in the Supplementary Table 1.

### Co-immunoprecipitation assay

For Co-IP in *Arabidopsis*, seeds were germinated and grown on ½MS. Seven-day-old seedlings were harvested and ground into powder with liquid nitrogen. The fine powder was resuspended in extraction buffer (50 mM Tris-HCl, pH 7.5, 150 mM NaCl, 0.5% [v/v] NP-40, and complete EDTA-free protease inhibitor cocktail [Roche]).

For Co-IP in tobacco, *Agrobacterium* cultures carrying the indicated constructs were resuspended in infiltration buffer (10 mM MgCl_2_, 10 mM methyl ethyl sulfide, pH 5.6, 100 µM acetosyringone) and incubated at room temperature for 2 h. The bacterial cultures were adjusted to a final OD_600_ = 0.5 for each construct and infiltrated into tobacco leaves. After 3 days incubation in the greenhouse, tobacco leaves were harvested, ground into powder with liquid nitrogen and then resuspended in extraction buffer (150 mM Tris-HCl, pH 7.5, 150 mM NaCl, 10% [v/v] glycerol, 10 mM EDTA, 1% [v/v] NP-40, and complete EDTA-free protease inhibitor cocktail [Roche]). The extracts were centrifuged at 20,000 × *g* at 4°C for 15 min. The supernatants were transferred to new 2-ml tubes and centrifuged for an additional 15 min. After centrifugation, the supernatants were incubated with GFP-trap magnetic agarose (Chromotek) at 4°C for 2 h. Beads were washed three times with 1 ml wash buffer (20 mM Tris-HCl, pH 7.5, 150 mM NaCl, 0.25% [v/v] NP-40] and then eluted with 80 µl elution buffer containing 1× NuPAGE™ LDS Sample Buffer (Thermo Fisher Scientific) and 1× NuPAGE® Reducing Agent (Thermo Fisher Scientific) and boiled for 10 min at 90°C.

### SDS-PAGE and Western Blotting

Total protein extracts were obtained by grinding the plant materials into a fine powder with liquid nitrogen. The powder was resuspended in extraction buffer (50 mM Tris-HCl, pH 7.5, 150 mM NaCl, 0.5% [v/v] NP-40, and complete EDTA-free protease inhibitor cocktail [Roche]). The extracts were centrifuged at 20,000 × *g* at 4°C for 15 min. The supernatants were transferred to new 2-ml tubes and centrifuged for an additional 15 min. The protein extracts were boiled in sample buffer (1× NuPAGE™ LDS sample buffer, 1× NuPAGE® Reducing Agent [Thermo Fisher Scientific]) for 10 min at 90°C and loaded on a 4–20% (w/v) Mini-PROTEAN® TGX™ Precast Protein Gels (Bio-Rad). The proteins were separated by electrophoresis and blotted onto a membrane with trans-Blot Turbo Mini 0.2 µm Nitrocellulose Transfer Packs (Bio-Rad). Membranes were blocked overnight at 4°C in 3% (w/v) skimmed milk dissolved in 25 mM Tris-HCl (pH 8), 150 mM NaCl and 0.05% (v/v) Tween20. The blots were then incubated at room temperature with the monoclonal α-GFP antibody (1:5000) (Miltenyi Biotec, 130-091-833), α-RFP antibody (1:5000) (ChromoTek, α-RFP antibody [6G6]), α-AP2S antibody^79^ (1:500), α-AP2A antibody^80^ (1:2000), α-AP1G antibody^81^ (1:2000), α-tubulin antibody (Sigma-Aldrich, T5168) (1:2000), α-CHC antibody (1:2000) (Santa Cruz, sc-57684) and α-TPLATE antibody (1:500)^82^.

### ICE analysis

The Sanger trace data of 9-day-old *p34^iCRISPR^-1* and *p34^iCRISPR^-2* plants were analysed with the online ICE Analysis Tool v3.0 (https://ice.synthego.com/). The percentage of genomic DNA sequences that contain an insertion or deletion in the sample is represented by indel %, whereas the knockout score corresponds with the proportion of sequences that will probably result in a functional protein knockout.

### Confocal microscopy and image quantification

All the confocal images, except Extended Data Fig. 7, were acquired with a Leica SP8X confocal microscope with application of the gating system (0.3 – 6 ns) for autofluorescence removal. GFP, YFP, RFP and FM4-64 were excited by white laser. The excitation and detection window settings were: GFP, 488/500-530 nm; YFP, 514/525-555 nm; RFP; 561/600-650 nm; and FM4-64; 515/570–670 nm. CFP was excited by diode at 405 nm and collected at 460-500 nm. Confocal images in Extended Data Fig. 7 were acquired with an inverted Olympus Fluoview1000 confocal by means of line sequential imaging at a resolution of 512×512 pixels, 8 µs/pixel scan speed, 4× line averaging, one-way scan direction, and an UPSLAPO 60x water immersion corrected lens (numerical aperture =1.20). The YFP and RFP channels were imaged with a 515-nm laser line from an argon laser (85% intensity) and a 559-nm solid state laser (10% intensity) and emission windows between 530 nm and 548 nm and between 580 nm and 615 nm, respectively. Confocal images in Fig. 6c,d and Extended Data Fig. 11b were acquired with a Zeiss LSM710 confocal laser scanning microscope equipped with a 40× water-corrected objective for detection of Alexa488 (488/500-545 nm), Alexa555 (561/555-610 nm) and DAPI (358/461 nm). BFA body size and BFA body number analyses (Fig. 6e) and the dark treatment for visualization of PIN2-GFP in vacuoles (Extended Data Fig. 11c) were measured as described previouly^83^. The fluorescent PM intensity and intracellular space were measured with ImageJ for the quantification of the FM4-64 uptake and the SYP61-CFP and VHAa-RFP signals. The relative fluorescent cytoplasm/PM intensity was calculated. The MTV1 signal was quantified by measurement of the skewness of the signal distribution as previously described^23^. Images were analysed by means of Fiji^84^ that was used to rotate and crop images.

### Statistical analysis

All statistical analyses were done with the GraphPad Prism v.8 software. The rBiFC data statistics were done with Brown-Forsythe and Welch analysis of variance (ANOVA) tests with Dunnett T3 multiple comparisons. For the statistical analysis of the protein levels, two-way ANOVA was utilized and followed by Sidak multiple comparison test in the comparison procedure. The other multiple comparisons were done by one-way ANOVA with Dunnett’s multiple comparisons test.

### Accession numbers

Sequence data from this article can be found in the Arabidopsis Genome initiative or GenBank/EMBL databases under the following accession numbers: *AP1G1* (AT1G23900), *AP1G2* (AT1G60070), *AP1S1* (AT2G17380), *AP2M* (AT5G46630), *AP2S* (AT1G47830), *TML* (AT5G57460), *AP3B* (AT3G55480), *AP4E* (AT1G31730), *AP4S* (AT2G19790), *AP5Z* (AT3G15160), *AP5B* (AT3G19870), *AP5M* (AT2G20790), *SPG11* (AT4G39420), *SPG15* (AT2G25730), *P34* (AT5G65960), *BAG4* (AT3G51780), *AAK1* (AT2G32850), *SH3P1* (AT1G31440), *SH3P2* (AT4G34660), *NECAP-1* (AT3G58600), *ECA1* (AT2G01600), *ECA4* (AT2G25430), *CAP1* (AT4G32285), *EPSIN2* (AT2G43160), *EPSIN3* (AT3G59290), *PICALM3* (AT5G35200), *AUXILIN-LIKE1* (AT4G12780), *AUXILIN-LIKE2* (AT4G12770), *VAMP721* (AT1G04750), *CLC1* (AT2G20760), *CLC2* (AT2G40060), *CHC1* (AT3G11130), *CHC2* (AT3G08530), *TOL6* (AT2G38410), *TOL9* (AT4G32760), *ACTIN2* (At3G18780), *BIN2* (AT4G18710), *KNOLLE* (AT1G08560) and *FH6* (AT5G67470).

## Supporting information

Proteomic analysis of adaptor complex interactome in Arabidopsis cell suspension cultures

Summary of protein-protein interactions tested by rBiFC assay.

## Acknowledgements

We thank I. Hwang, G. Jürgens and J. Pan for the kind gift of the α-AP1G and α-AP2A, the α-KNOLLE and α-AP2S antibodies, respectively, M. Sauer, S. Schneider, J. Friml, J. Lin and I. Hara-Nishimura for providing published materials, T. Jacobs for useful discussions and M. De Cock for help in preparing the manuscript. This work was supported by the Research Foundation-Flanders projects (G008416N, G0E5718N and 3G038020 to E.R.), the Belgian Science Policy Office for a postdoctoral fellowship (R.K.), the China Scholarship Council for predoctoral fellowships (P.W., X.Z. and R.W.), and the European Research Council T-Rex (project number 682436 to D.V.D.).

## Author contributions

W.S., P.W. X.Z. and E.R. initiated the project and designed experiments. W.S., X.Z., R.K., A.H. and N.D.W. did cloning for TAP-MS and AP-MS. D.E., J.V.L., K.G. and G.D.J performed the MS work and analysed data. W.S. and P.W. did the interactome validation. P.W. performed all the P34 work. E.M. did microscopy. R.A.K. and C.T. contributed materials. D.A., M.V. and D.V.D did the proximity labelling. R.W and S.V. performed the PIN2 immunolabelling. W.S., P.W. and E.R. wrote the manuscript. All authors revised the manuscript.

## Additional information

### Supplementary information

The online version contains supplementary material

Correspondence and requests for materials should be addressed to W.S. and E.R.

**Extended Data Fig. 1.**
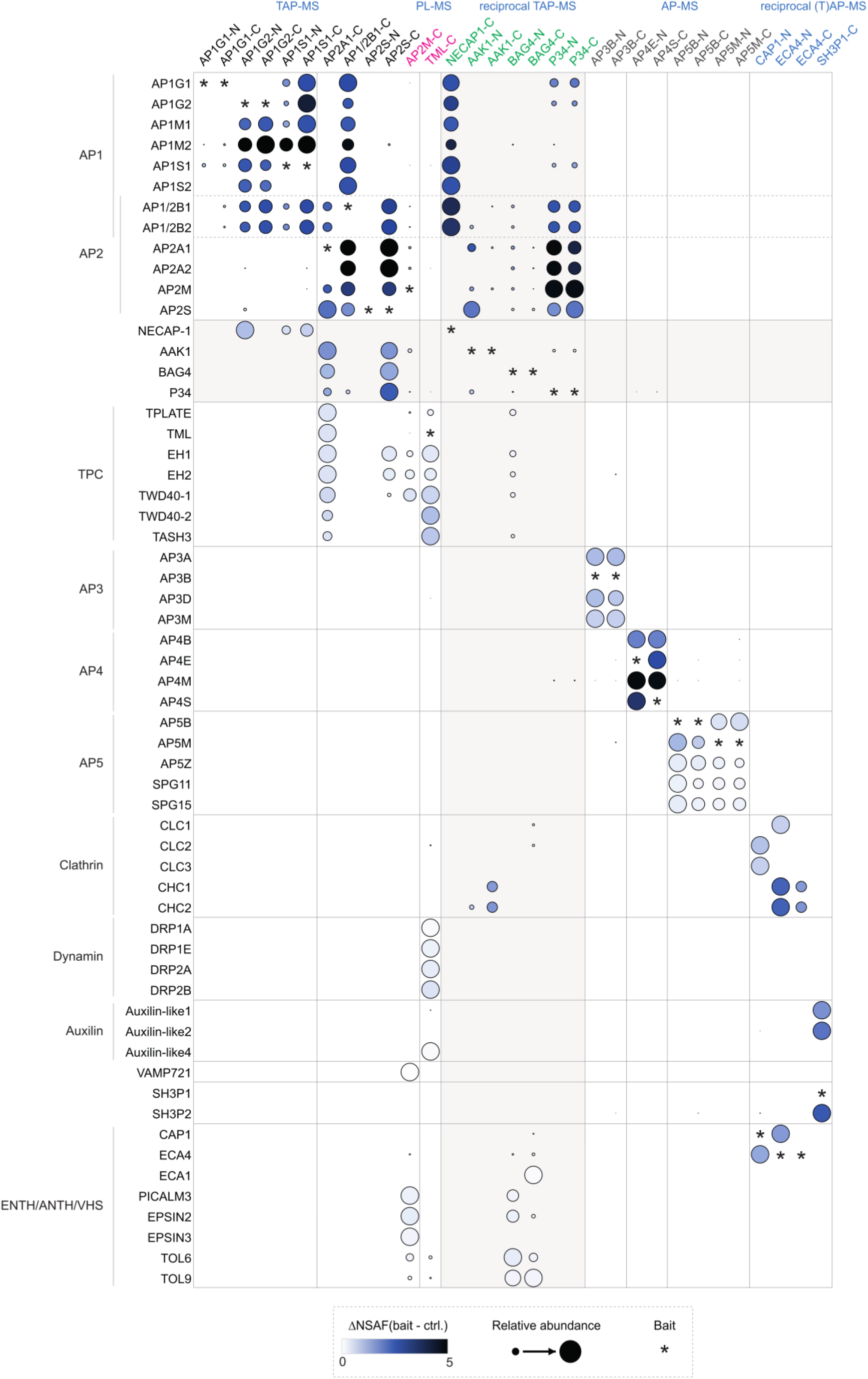
Dot plot matrix of selected proteins from the AP/TPC interactome. Quantitative dot plot matrix covering core AP/TPC subunits and a selection of proteins linked to endocytosis/vesicle trafficking. The colour hue of the nodes corresponds with the abundance of each prey in a given experiment, calculated by subtracting the average normalized spectral abundance factor (NSAF) in the control dataset from the average NSAF (bait) of each prey [NSAF (bait - ctrl.)]. The size of the dots reflects the relative abundance of each prey over the different experiments. The identification of each bait protein is shown by an asterisk.

**Extended Data Fig. 2.**
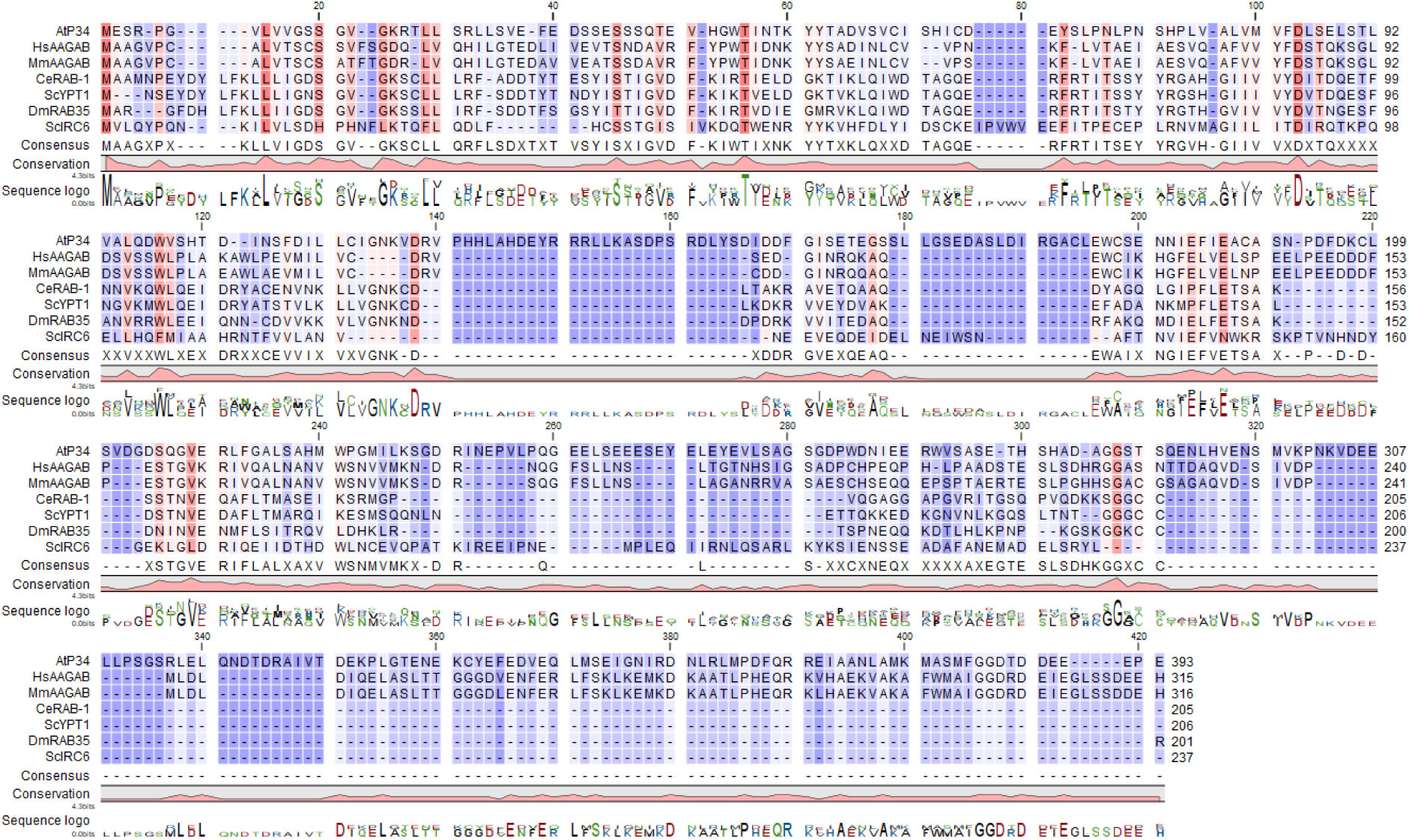
Protein sequence alignment of P34 in eukaryotes. Amino acid sequence alignment of the P34/AAGAB family in At, *Arabidopsis thaliana*; Hs, *Homo sapiens*; Mm, *Mus musculus*; Ce, *Caenorhabditis elegans*; Sc, *Saccharomyces cerevisiae*; Dm, *Drosophila melanogaster*. The sequences were aligned with the CLC Main Workbench (Qiagen). The colour intensity reflects how conserved a particular position is in the alignment. Dark orange and dark purple represents 100% and 0% identity, respectively.

**Extended Data Fig. 3.**
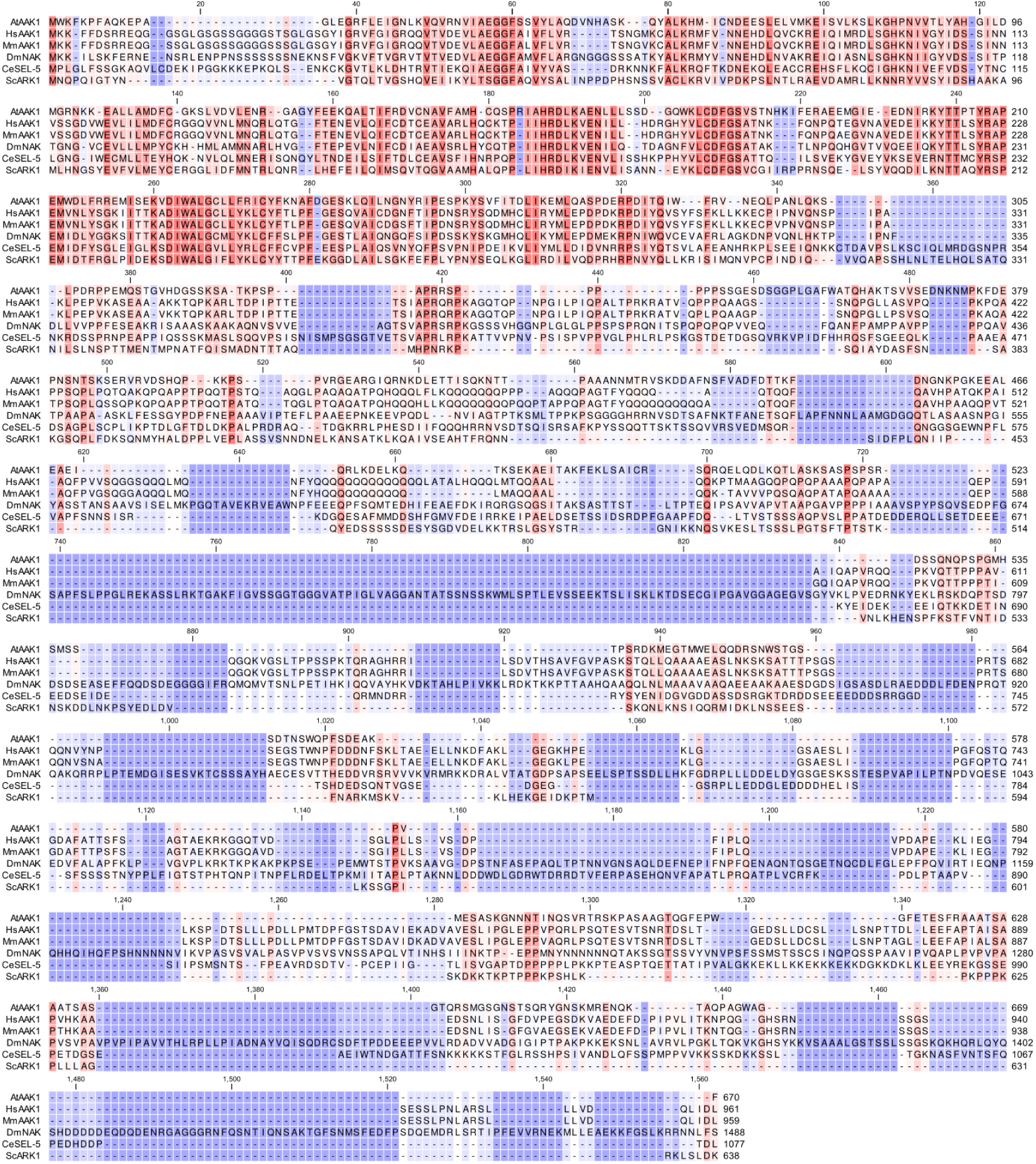
Protein sequence alignment of AAK1 in eukaryotes. Amino acid sequence alignment of the NAK family in At, *Arabidopsis thaliana*; Hs, *Homo sapiens*; Mm, *Mus musculus*; Dm, *Drosophila melanogaster*; Ce, *Caenorhabditis elegans*, and Sc, *Saccharomyces cerevisiae*. The sequences were aligned with CLC Main Workbench (Qiagen). The colour intensity reflects how conserved the particular position is in the alignment. Dark orange and dark purple represent 100% and 0% identity, respectively.

**Extended Data Fig. 4.**
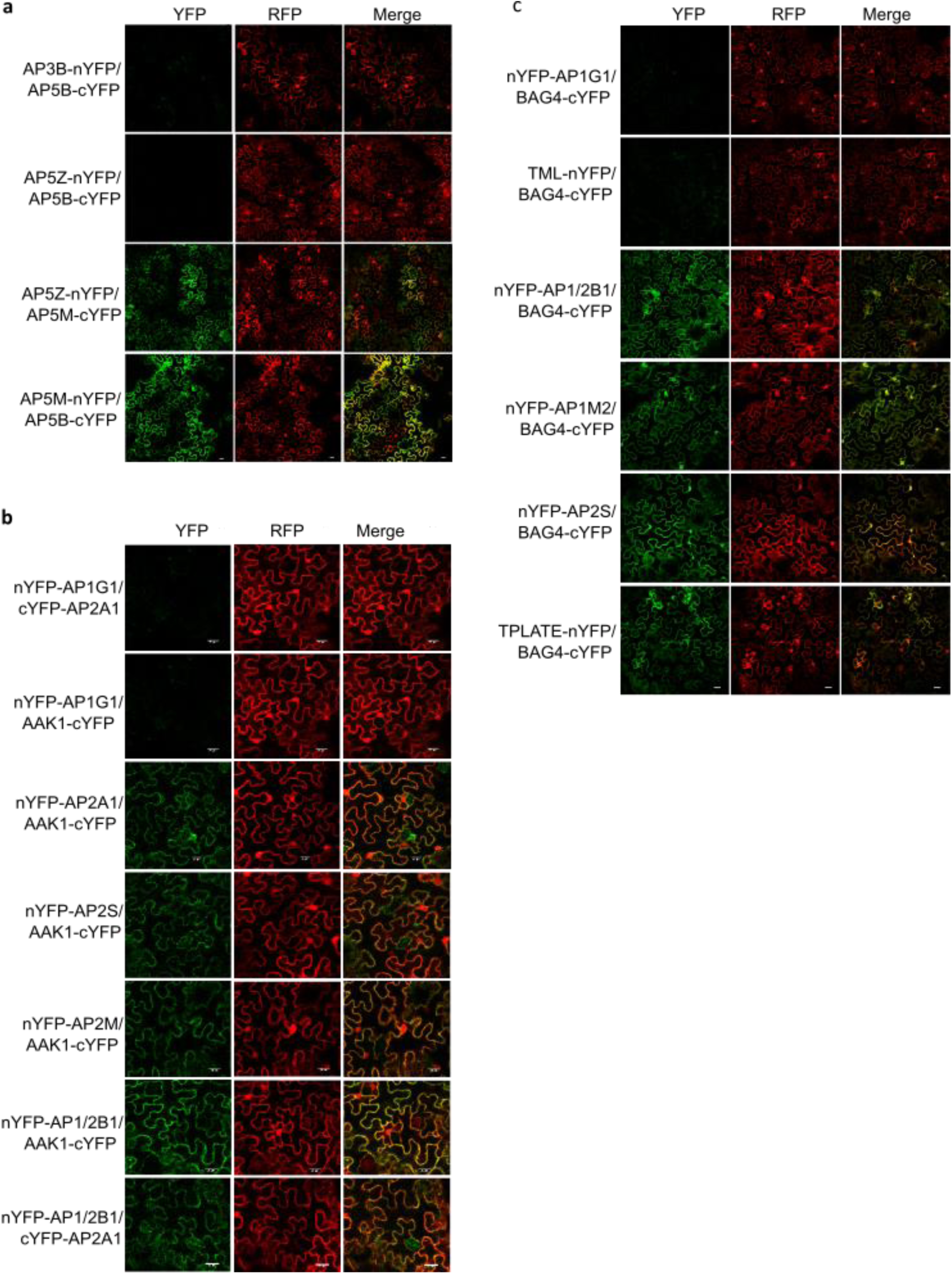
rBiFC analysis with AP-5, AAK1 and BAG4. Confocal images of the rBiFC assay of AP-5 (a), AP-2 (b) and BAG4 (c) subunits with different AP subunits quantified in Figures 4b, 4e and 4h, respectively. Scale bars, 20 µm.

**Extended Data Fig. 5.**
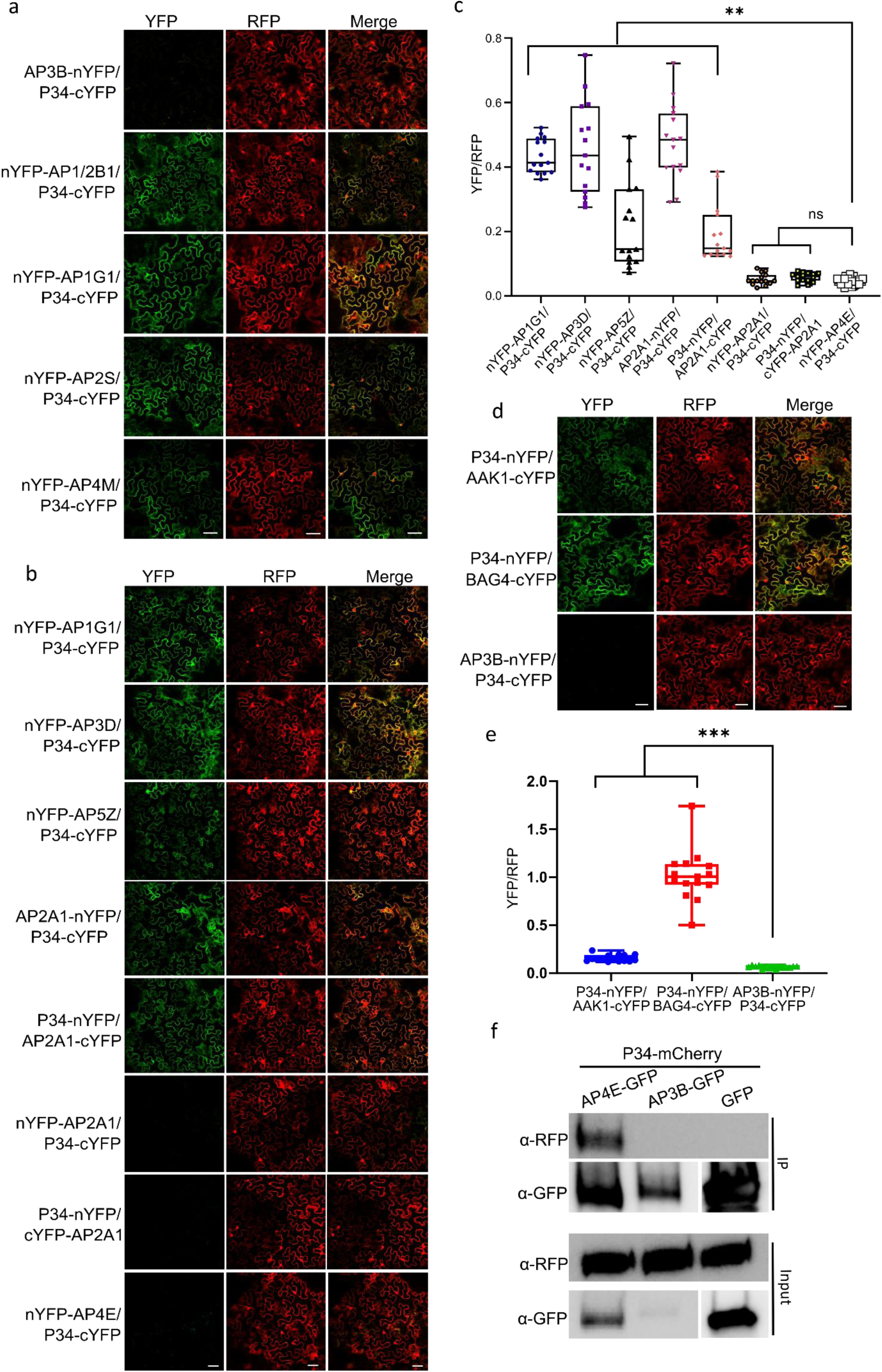
rBiFC and Co-IP analyses with P34. **a**, **b,** and **d**, Confocal images of the rBiFC assays of P34 with different AP subunits (quantified in Figure 4k), with AAK1 and with BAG4, respectively. **c** and **e,** Quantification of the rBiFC (YFP/RFP ratio) in (b) and (d) respectively. *n*=15, *n,* number of cells analysed. \*\**P* ≤ 0.01, \*\*\**P* ≤ 0.001 (Brown-Forsythe and Welch ANOVA tests combined with Dunnett T3 multiple comparisons test); ns, not significant. **f,** Validation of the interactions between P34, AP3B, and AP4E by co-immunoprecipitation (Co-IP) in tobacco leaves transiently expressing the *p35S:P34:mCherry*, *p35S:AP3B-GFP* and *p35S:AP4E-GFP* constructs. The *p35S:GFP* construct was used as a negative control. rBiFC experiments were repeated twice and one representative experiment is shown. Scale bars, 50 µm.

**Extended Data Fig. 6.**
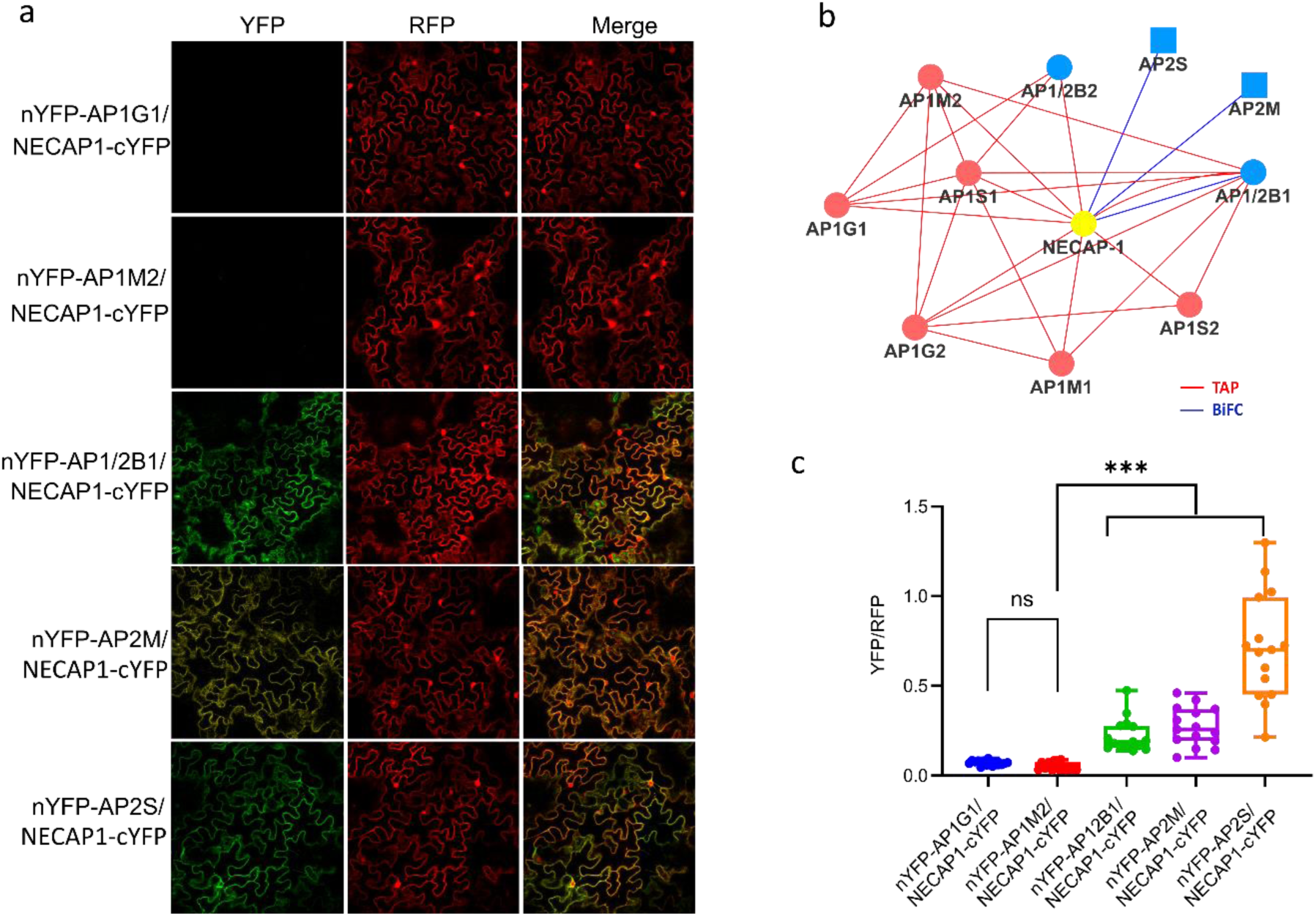
Validation of the interactions with NECAP-1 by rBiFC. **a**, Confocal images of the rBiFC assay of NECAP-1 with different AP-1 and AP-2 subunits. Scale bars, 20 µm. **b**, Cytoscape model summarizing the interactions between various subunits and NECAP-1. Edge colours indicate the analysis method. Node colours correspond to the different complexes and protein families, red and blue, for AP-1 and AP-2, respectively. **c**, Quantification of rBiFC (YFP/RFP ratio) in (a). rBiFC experiments were repeated twice and one representative experiment is shown. *n*=15, *n,* number of cells analysed. \*\*\**P* ≤ 0.001 (Brown-Forsythe and Welch ANOVA tests combined with Dunnett T3 multiple comparisons test); ns, not significant.

**Extended Data Fig. 7.**
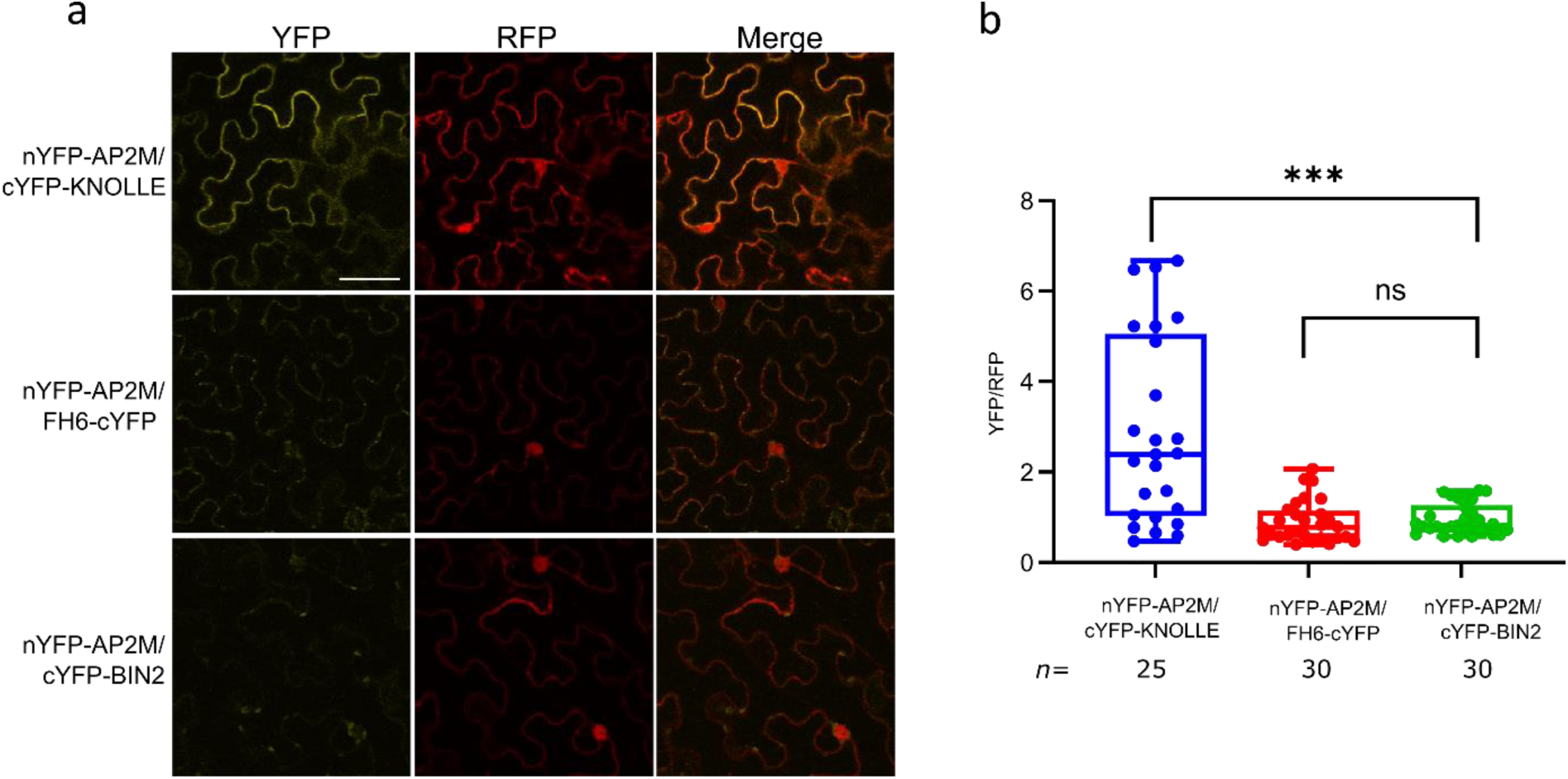
BiFC confirmation of the AP2M cargo interactions discovered by the proximity labelling (PL) experiments. **a,** Confocal images of the rBIFC assay of AP2M with the two interactors, KNOLLE and FORMIN-LIKE PROTEIN 6 (FH6), observed by PL-MS with AP2M as bait. The SHAGGY-like kinase BIN2 was used as negative control. Whereas AP2M interacts with KNOLLE evenly along the plasma membrane, the interaction between AP2M and FH6 is less intense and concentrated in plasma membrane-associated punctate. AP2M and BIN2 do not visually interact and only background and chlorophyll fluorescence are observed. **b,** Quantification of the rBiFC (YFP/RFP PM intensity ratio) in (a) clearly showing significant differences between AP2M-KNOLLE and AP2M-BIN2. The punctate signal in the AP2M-FH6 combination is not sufficiently strong to yield a statistical difference compared to the control. *n,* number of cells analysed. *\*\*\*P* < 0.001 (Brown-Forsythe and Welch ANOVA tests with Dunnett T3 multiple comparisons test); ns, not significant. Scale bar, 50 µm.

**Extended Data Fig. 8.**
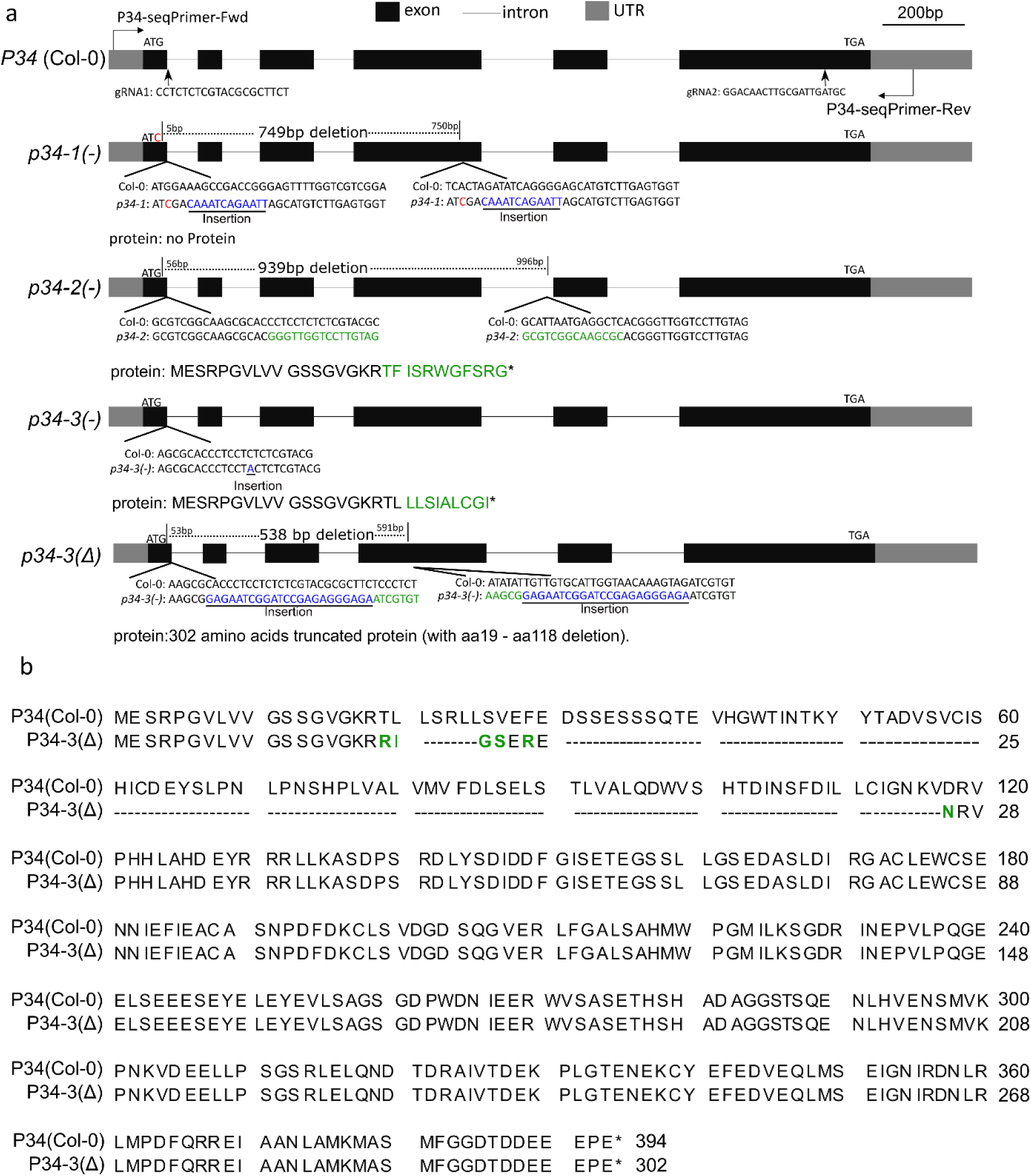
Genotype of the *P34-CRISPR* lines. **a,** Schematic representation of the CRISPR editing on the *P34* gene in the *p34* CRISPR mutants. Blue, red, and green indicate insertion, nucleotide exchange, and mutated amino acids, respectively. The arrows mark the sites of the guide RNA sequences (gRNA). **b,** Protein sequence alignment of P34 (wild type, Col-0) and P34(Δ) by means of the CLC Main Workbench (Qiagen). Hyphen and green letters indicate deletion and mutated amino acids, respectively. Asterisks indicate the stop codons.

**Extended Data Fig. 9.**
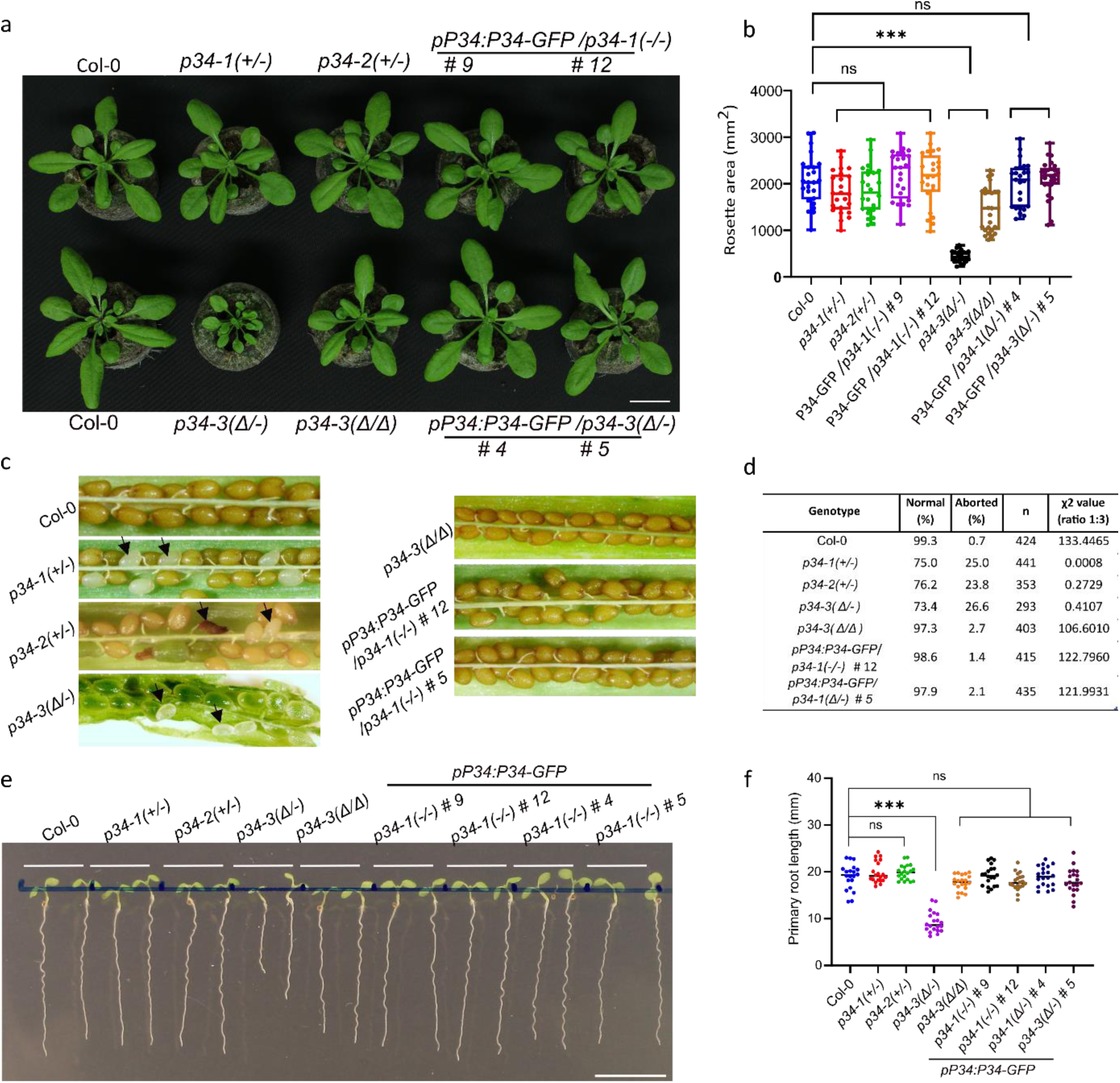
Phenotype of the *P34-CRISPR* lines. **a**, Rosettes of the wild type (Col-0), the *p34-1(+/-), p34-2(+/-), p34-3(Δ/-), p34-3(Δ/Δ)* mutants and the *p34-1(-/-)* (lines #9 and #12) and *p34-3(Δ/-)* (lines #4 and #5) mutants complemented with the *pP34:gP34-GFP* constructs and grown in the soil for 4 weeks. Scale bar, 20 mm. **b**, Quantification the rosette leaf area of each genotype show in (a). Three independent experiments were combined, each with 8-9 plants per genotype in each experiment. \*\*\**P* ≤ 0.001 (one-way ANOVA test); ns, not significant. **c,** Aborted seeds in the siliques of genotypes shown in (b). Arrows indicate the aborted seeds. **d**, Quantification of the aborted seeds in (**c**). χ^2^ values were calculated with the χ^2^ test. *n* number of seeds analysed. **e**, Phenotype of seven-day-old seedlings grown on ½MS. Scale bar,10 mm. **f**, Primary root length of seedlings shown in (f). 20 roots per genotype were measured. \*\*\**P* ≤ 0.001 (one-way ANOVA test).

**Extended Data Fig. 10.**
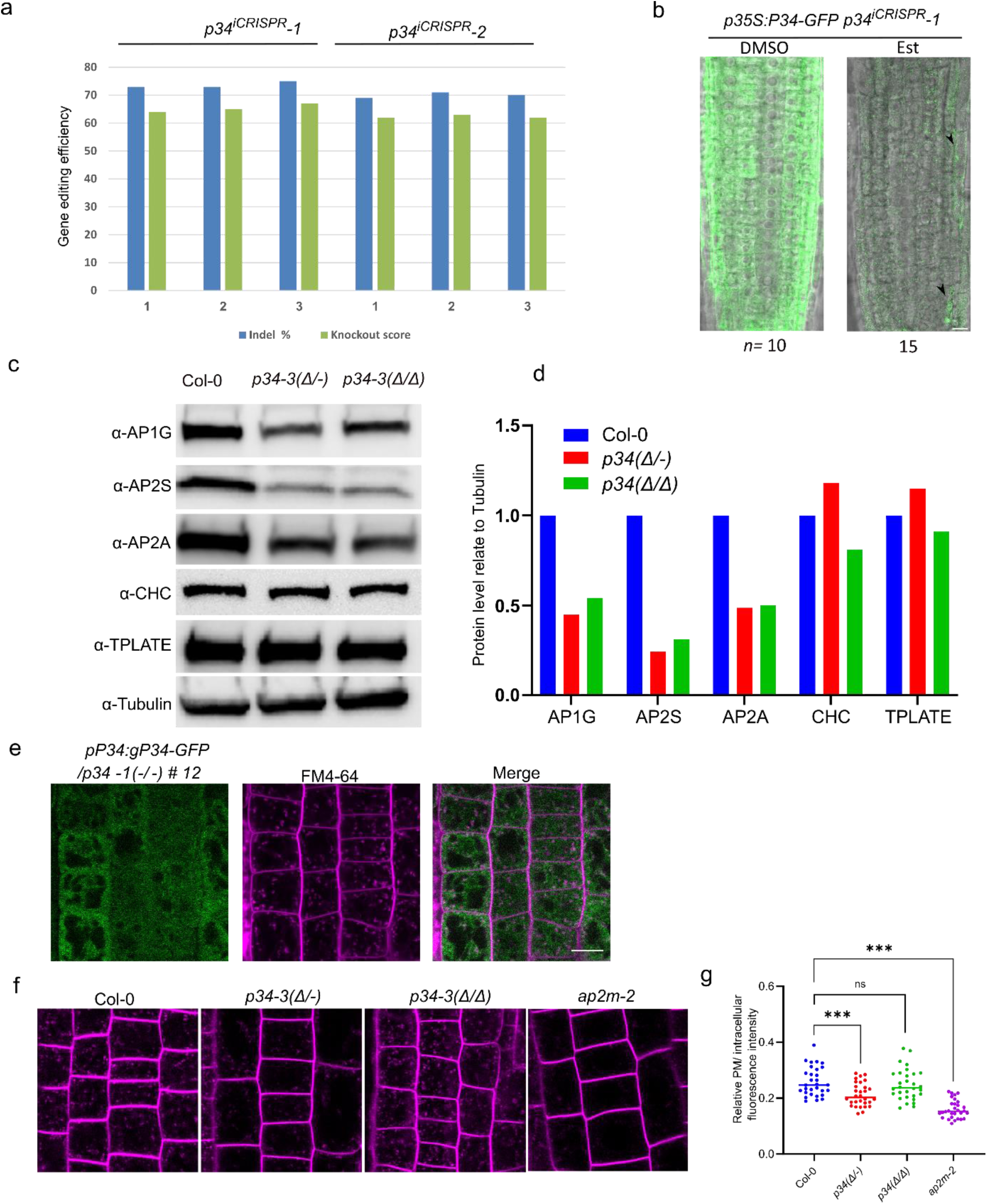
CRISPR efficiency and different phenotypes of the *p34* mutants. **a**, Gene editing efficiency analysis of three random 9-day-old plants of *p34^iCRISPR^-1* and *p34^iCRISPR^-2* after β-estradiol induction. The sequencing results were analysed with the online software Inference of CRISPR Edits (ICE) (https://ice.synthego.com/<u>#/</u>). **b**, P34-GFP localization in the roots of 5-day-old *p35S:gP34-GFP/p34^iCRISPR^-1* plants grown on DMSO and 10 μM β-estradiol (Est). Arrows indicate the remaining P34-GFP signal. Scale bar, 10 µm. **c,** Protein levels of AP1G, AP2A and AP2S in *p34-3* mutants analysed by immunoblotting with α-AP1G, α-AP2S, α-AP2A, α-CHC, α-TPLATE and α-Tubulin. **d,** Quantification of protein levels shown in (c). The protein level was normalized to tubulin. **e,** Confocal images of the 5-day-old *pP34:gP34-GFP/p34-1(-/-)* (line *#*12) root cells stained with FM4-64 (2 μM, 40 min). Scale bar, 10 µm. **f,** FM4-64 uptake in the *p34-3* mutants. **g**, Fluorescence intensity ratio of the relative intracellular-to-plasma membrane (PM) FM4-64 signal. \*\*\**P* ≤ 0.001 (one-way ANOVA test); ns, not significant. *n*=30, *n* number of cells analysed, Scale bar, 10 µm.

**Extended Data Fig. 11.**
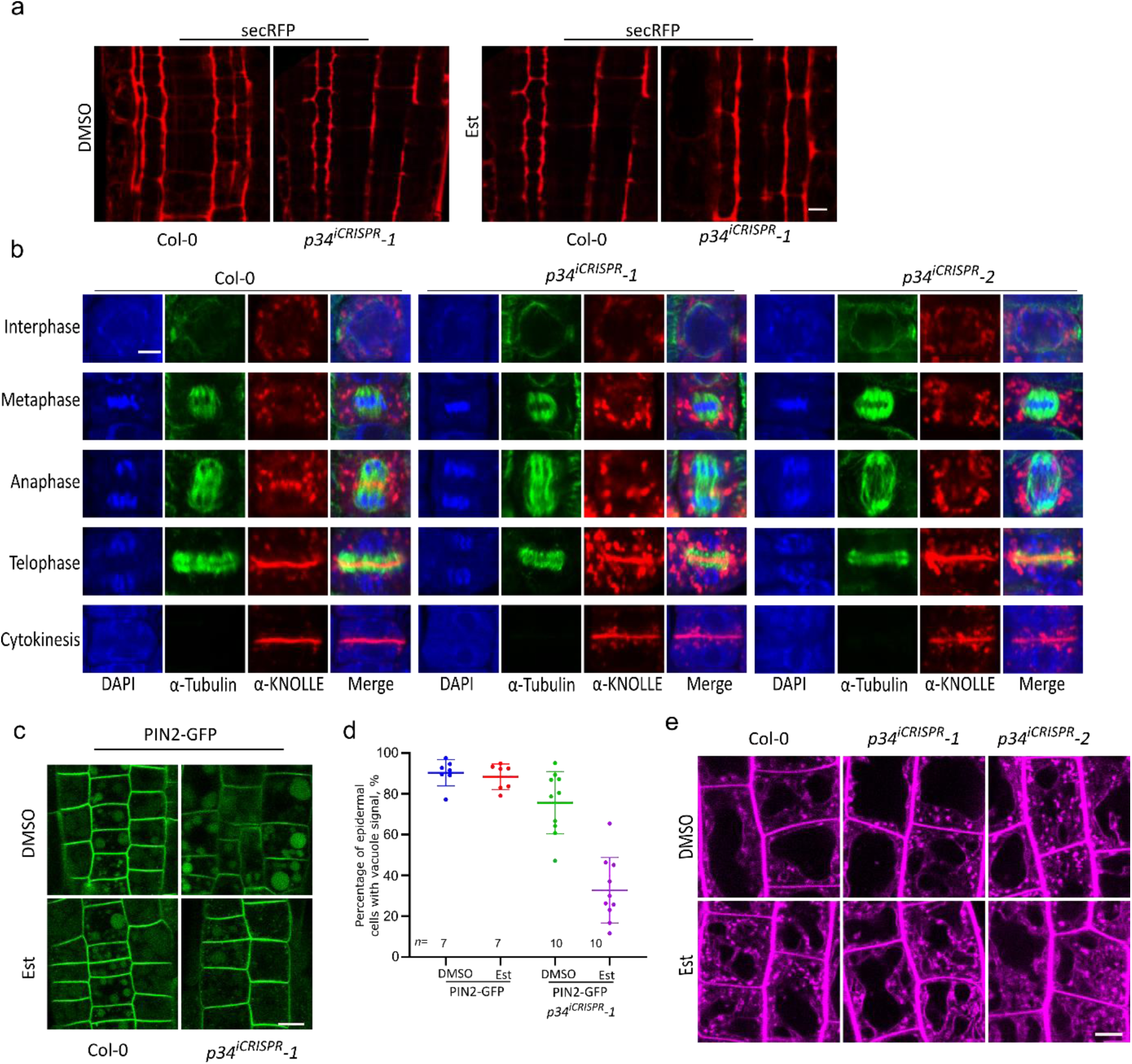
Vesicular trafficking pathways affected in the *p34* mutants. **a**, secRFP localization in roots of 5-day-old Col-0 and *p34^iCRISPR^-1* plants grown on DMSO and 10 μM β-estradiol (Est). **b**, Immunofluorescence staining of KNOLLE in the root meristem of Col-0, *p34^iCRISPR^-1* and *p34^iCRISPR^-2* plants treated with 10 μM β-estradiol. **c**, PIN2-GFP localization in the roots of 5-day-old Col-0 and *p34^iCRISPR^-1* plants grown on DMSO and 10 μM β-estradiol in the dark. **d,** Percentage of epidermal cells with GFP signals in the vacuoles as shown in (c). **e**, Morphology of the lytic vacuoles in root cells of 7-day-old Col-0, *p34^iCRISPR^-1* and *p34^iCRISPR^-2* plants grown on DMSO and 10 μM β-estradiol. Images were taken after 2 h of staining with 4 μM FM4-64. Scale bars, 10 µm (a-c, e)

**Supplementary Table 1.**
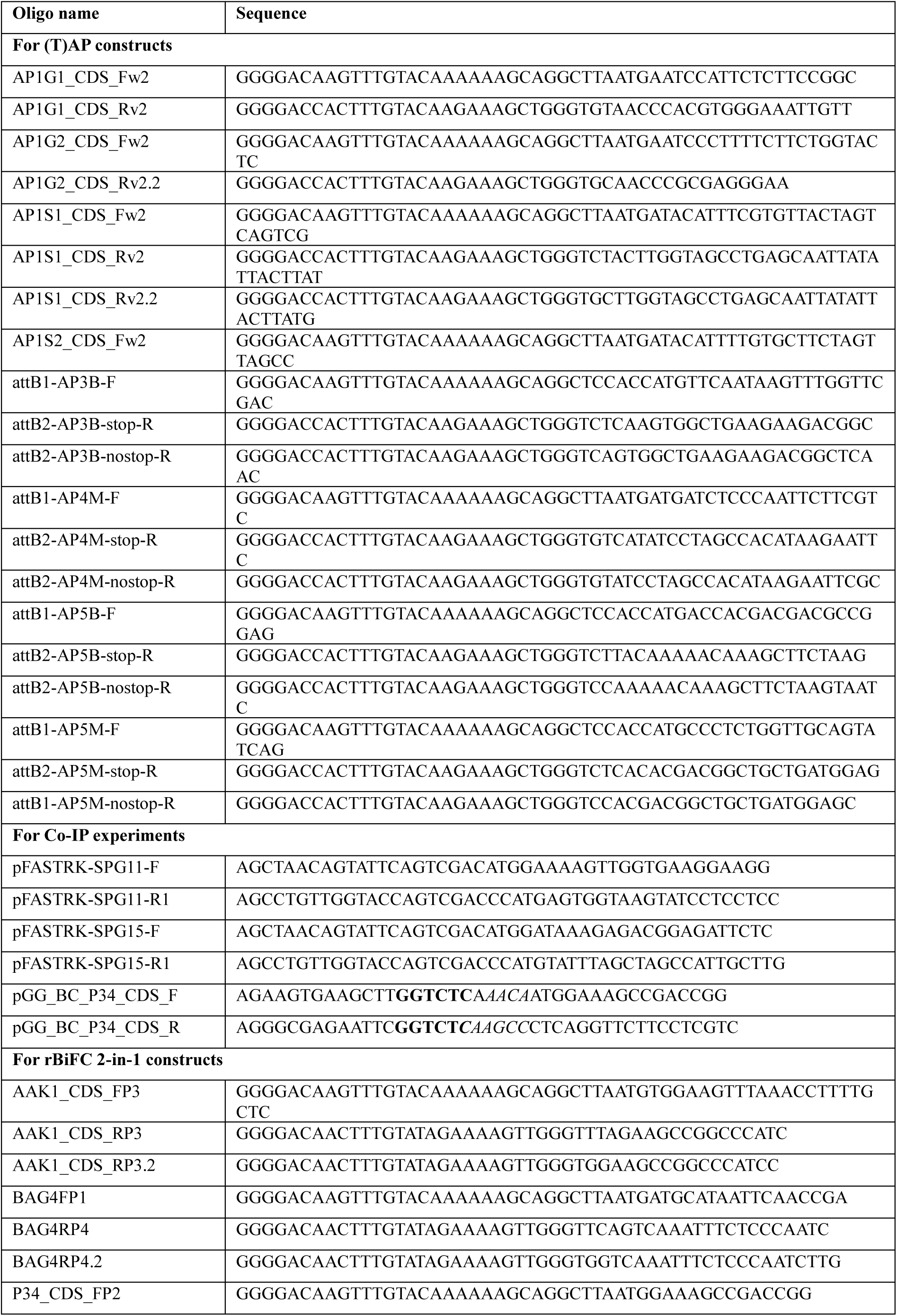
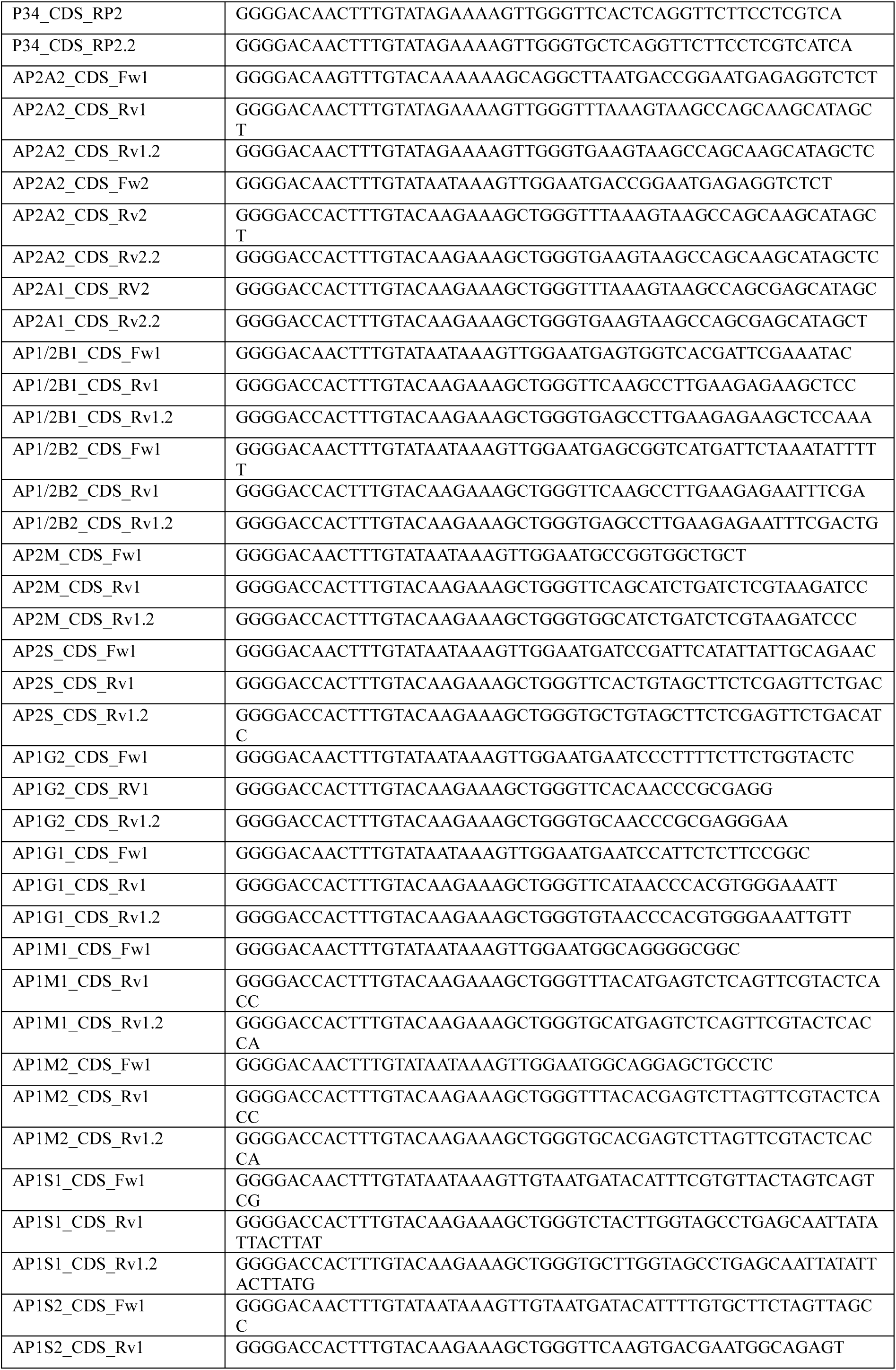
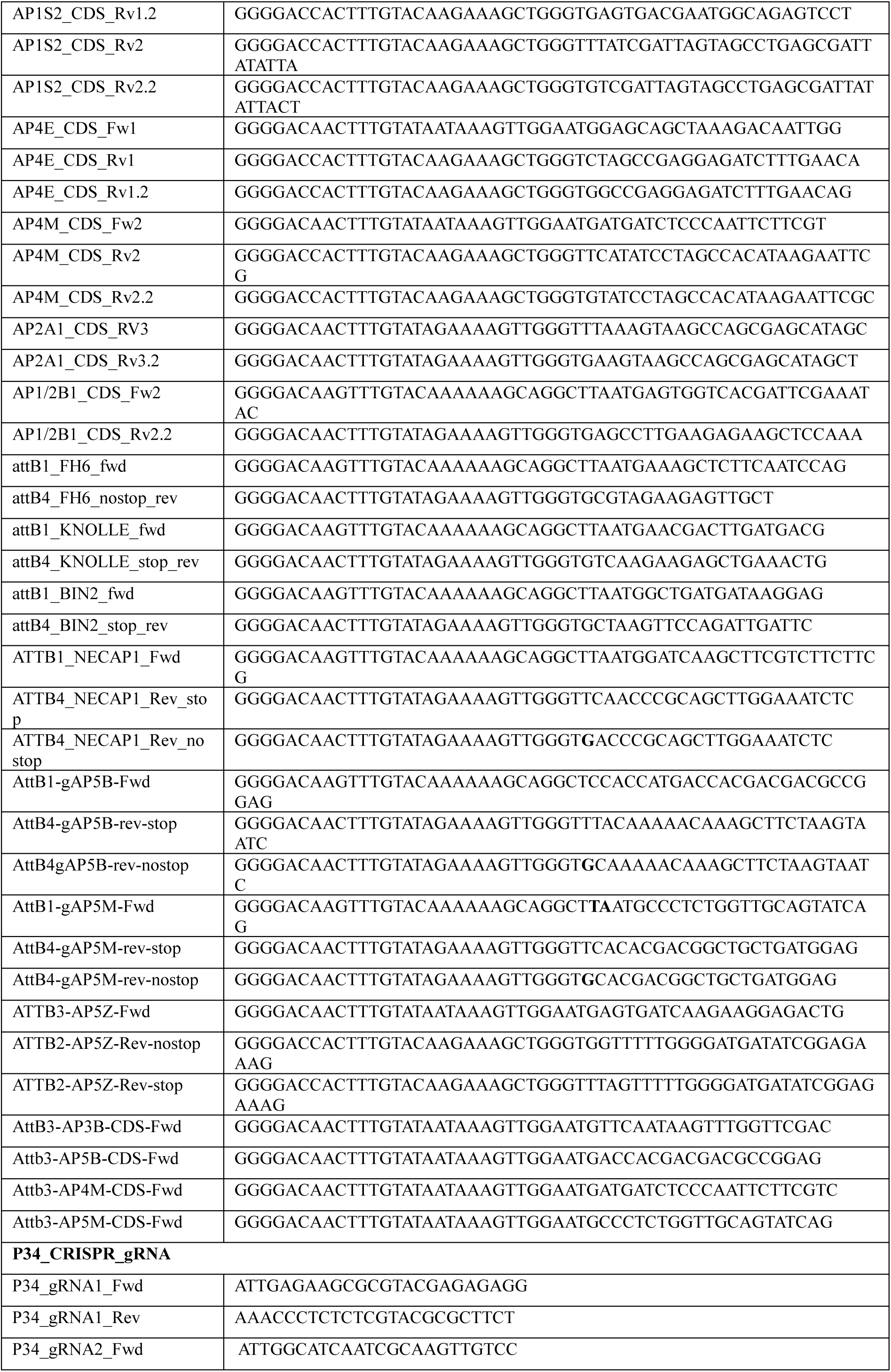
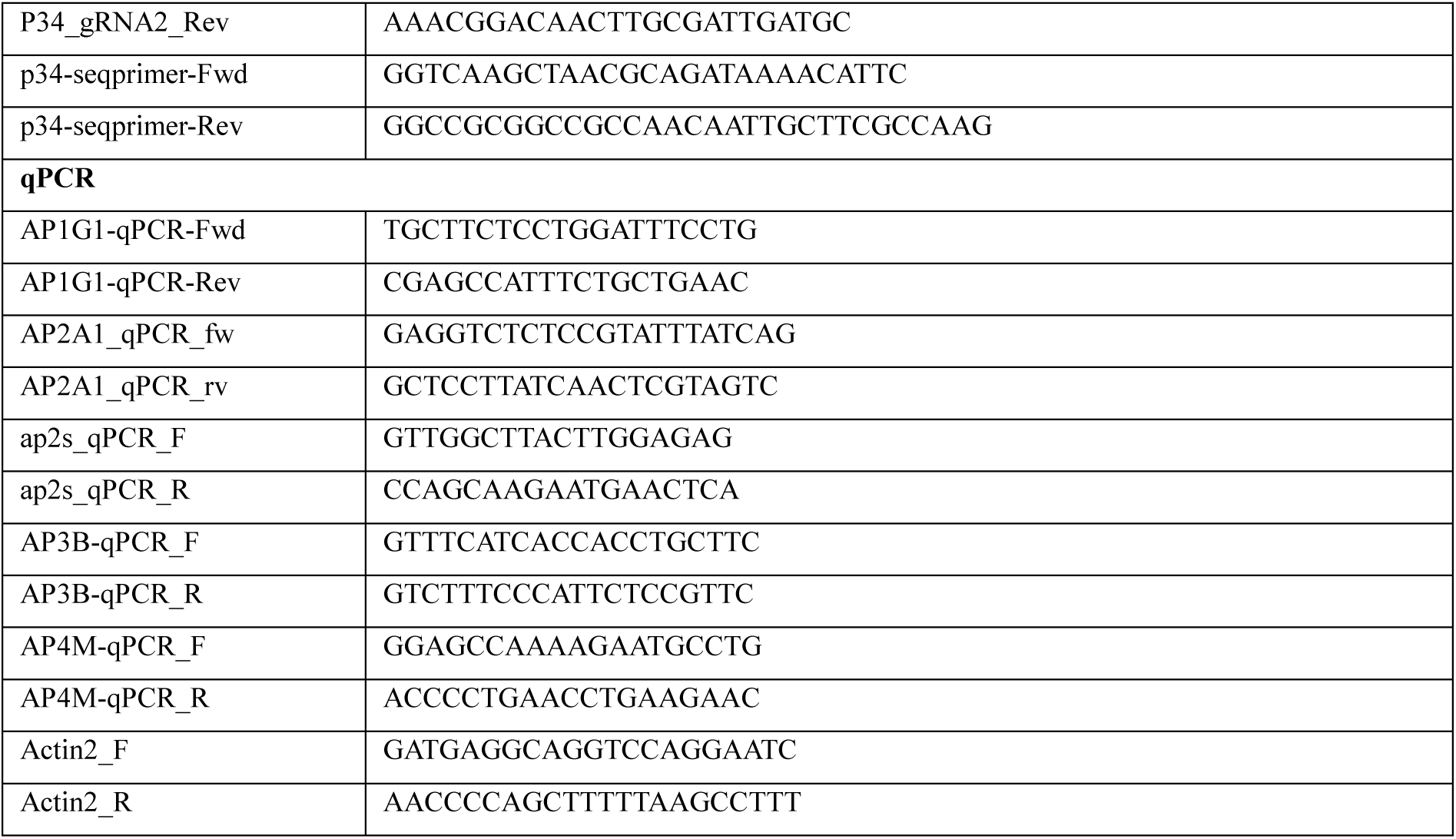
Oligonucleotides used in this study.

